# Cosmopolitanism at the Roman Danubian Frontier, Slavic Migrations, and the Genomic Formation of Modern Balkan Peoples

**DOI:** 10.1101/2021.08.30.458211

**Authors:** Iñigo Olalde, Pablo Carrión, Ilija Mikić, Nadin Rohland, Shop Mallick, Iosif Lazaridis, Miomir Korać, Snežana Golubović, Sofija Petković, Nataša Miladinović-Radmilović, Dragana Vulović, Kristin Stewardson, Ann Marie Lawson, Fatma Zalzala, Kim Callan, Željko Tomanović, Dušan Keckarević, Miodrag Grbić, Carles Lalueza-Fox, David Reich

## Abstract

The Roman Empire expanded through the Mediterranean shores and brought human mobility and cosmopolitanism across this inland sea to an unprecedented scale. However, if this was also common at the Empire frontiers remains undetermined. The Balkans and Danube River were of strategic importance for the Romans acting as an East-West connection and as a defense line against “barbarian” tribes. We generated genome-wide data from 70 ancient individuals from present-day Serbia dated to the first millennium CE; including Viminacium, capital of Moesia Superior province. Our analyses reveal large scale-movements from Anatolia during Imperial rule, similar to the pattern observed in Rome, and cases of individual mobility from as far as East Africa. Between ∼250-500 CE, we detect gene-flow from Central/Northern Europe harboring admixtures of Iron Age steppe groups. Tenth-century CE individuals harbored North-Eastern European-related ancestry likely associated to Slavic-speakers, which contributed >20% of the ancestry of today’s Balkan people.

## Introduction

During the 2^nd^ century CE, the Roman Empire stretched from Mesopotamia in the East to Iberia in the West, controlling the entirety of the Mediterranean Sea and bordered in the North by the Rhine and Danube rivers. Despite being connected through the world’s first systematic road network and rapid sea links, human populations inhabiting this vast area were extremely diverse culturally and also genetically.

The ancestry make-up of the different regions of the Roman Empire have not been systematically characterized. Among seven Romans from York (ancient Eboracum), in England, one individual showed affinities to modern Middle East populations^1^, and individuals with a high proportion of North-African ancestry were found in Roman southern Iberia ^2^. Analysis of 48 skeletons from the Roman Imperial period in the Lazio region of Italy (the region of the capital) showed that at the height of the Empire, genetic ancestry become much more heterogeneous than in previous periods and shifted towards Near Eastern populations ^3^. These results suggest a considerable degree of diversity across the Empire during the Imperial period^4^. A key region of the Roman Empire was the Danubian *limes* (the Roman military frontier zone) which to a large extent falls in the territory of present-day Serbia, and which was indubitably one of the most strategically important border areas. Throughout history, the Balkan peninsula has been a crossroad of different cultures due to its geographical location between East and West. During the Imperial period, this region was politically central as reflected in the fact that 18 Roman Emperors were born in the territory of modern Serbia, including Constantine the Great. The Roman Balkans was divided into two areas of cultural influence separated by the so-called Jireček line ^5^: the south was mostly influenced by Greek language and culture whilst the north was mostly Latin influenced. During the Migration Period, the Balkans became a channel for movements of people towards the west and the south: Goths, Huns, Gepids, Heruli, Lombards or Slavs moved through, sacked and/or settled in the Balkan territories ^6^. The Slavs, who raided the Balkans during the 6th century and settled in the region in the 7th century ^6^, had a particularly long-lasting cultural impact, reflected in the Slavic languages widely spoken in the region today including Serbian, Croatian, Montenegrin, Bosnian, Slovenian, Macedonian, and Bulgarian. Even present-day Peloponnesians (southern tip of the Balkan Peninsula) carry a small yet significant amount of Slavic-related ancestry ^7^, but the ultimate origin of Slavic speakers and the degree of demographic impact in the region is not yet well understood.

We generated genome-wide data from 70 ancient individuals from 3 archaeological sites in present-day Serbia, dating from ∼250-500 CE and 900-1000 CE. To place the results in a geographic and temporal context, we also generated genome-wide data from present-day Serbs and analyzed them alongside other Balkan groups.

## Results

### Data generation

We built double-stranded DNA libraries with partial uracil-DNA glycosylase (UDG) treatment and used in-solution hybridization ‘1240k’ enrichment ^8,9^ to generate genome-wide data as well as mitochondrial DNA sequence data. We successfully obtained data from 72 individuals collected from 3 archeological sites in present-day Serbia dating from the 1^st^ millennium CE (Supplementary TABLE 1). We also used the Affymetrix Human Origins array ^10^ to generate genome-wide data from present-day Serb individuals from throughout the Balkans (*n*=37) (see Supplementary section 1).

A total of 52 individuals for whom we obtained data were excavated from the Roman city and military encampment of Viminacium, which was located at the confluence of the Mlava River and the Danube and was the capital of Upper Moesia province; the individuals came from the four necropoli of Pirivoj (*n*=19), Vise Grobalja (*n*=10), Rit (*n*=13) and Pećine (*n*=10) (Supplementary section 2.1) ^11^. Another 18 individuals were excavated from Timacum Minus probably which consisted of a military fortification and adjacent village on the left bank of the Beli Timok River and probably acted as the administrative hub of regional mining operations ^12^; the individuals fcame from the two necropoli of Slog (*n*=10) and Kuline (*n*=7). Finally, ^12^two individuals came from the Mediana archeological site, as a *suburbium* of the neighboring city of Naissus, located near the Roman *via publica* which connected the major urban centers of the Dacia diocese such as Viminacium, Singidunum and Naissus ^13^ (Figure S1B).

We generated 20 radiocarbon dates (Table ST1), which confirmed that the necropoli dated to ∼1-500 CE, with the exception of Kuline at ∼900-1000 CE. We discarded 11 individuals with fewer than 20,000 SNPs from genome-wide analyses. Of the remaining 59, only one (I15503) had weak evidence of contamination based on the rate of mismatch in the mtDNA genome and ratio of Y-chromosome to X+Y-chromosome reads and was flagged as a questionable source of genetic information (Supplementary TABLE 1). We identified two close kinship relationships (Supplementary section 7): a pair of identical twins (I15538 and I15539) from Kuline necropolis (Timacum Minus) found in two adjacent graves, and a pair of second-degree relatives (I15490 and I15491) from a double grave in Pirivoj Necropolis (Viminacium).

### Extraordinary genetic heterogeneity

We performed Principal Component Analysis (PCA) by projecting ancient individuals onto the axes computed on a set of present-day West-Eurasian (WE) populations (Figure S7); and onto a set of present-day West-Eurasian (WE), North African (NA) and Sub-Saharan populations (Figure S8), all genotyped on the Human Origins array.

Ancient individuals from Viminacium, Timacum Minus and Mediana plotted diffusely, ranging from Near Eastern populations on the right of PC1 to European populations on the left (Figure S7). One individual (I15499) from Viminacium projected outside West Eurasian genomic diversity. Heterogeneous ancestry is also inferred in ADMIXTURE analysis (Figure S10). A key feature of the data is two parallel genetic clines running along PC1 (Fig. 1). We call the first the “ Balkan Iron Age cline”, with southern Balkan populations such as Bronze Age and Iron Age Aegean groups on the right extreme closer to Near Eastern populations (larger values in PC1), northern populations such as Slovenian Iron Age groups on the left extreme closer to Central European populations (smaller values in PC1), and a Bulgarian Iron Age individual and Bronze Age and Iron Age Croatian groups taking intermediate positions but closer to the southern and northern extremes, respectively. This Iron Age cline is mirrored by the “ present-day Balkan cline”, which is shifted towards the upper-left of the plot (lower values in PC1 and higher values in PC2) with respect to the Iron Age cline but maintains the same geographical pattern of southern Balkan populations such as the Greeks on the right, and northern Balkan populations such as Croatians on the left. This suggests that present-day populations are not direct descendants without admixture of Iron Age groups from the same region, and that similar demographic shaped Balkan populations from North to South over the past 2,000 years.

**Figure 1.**
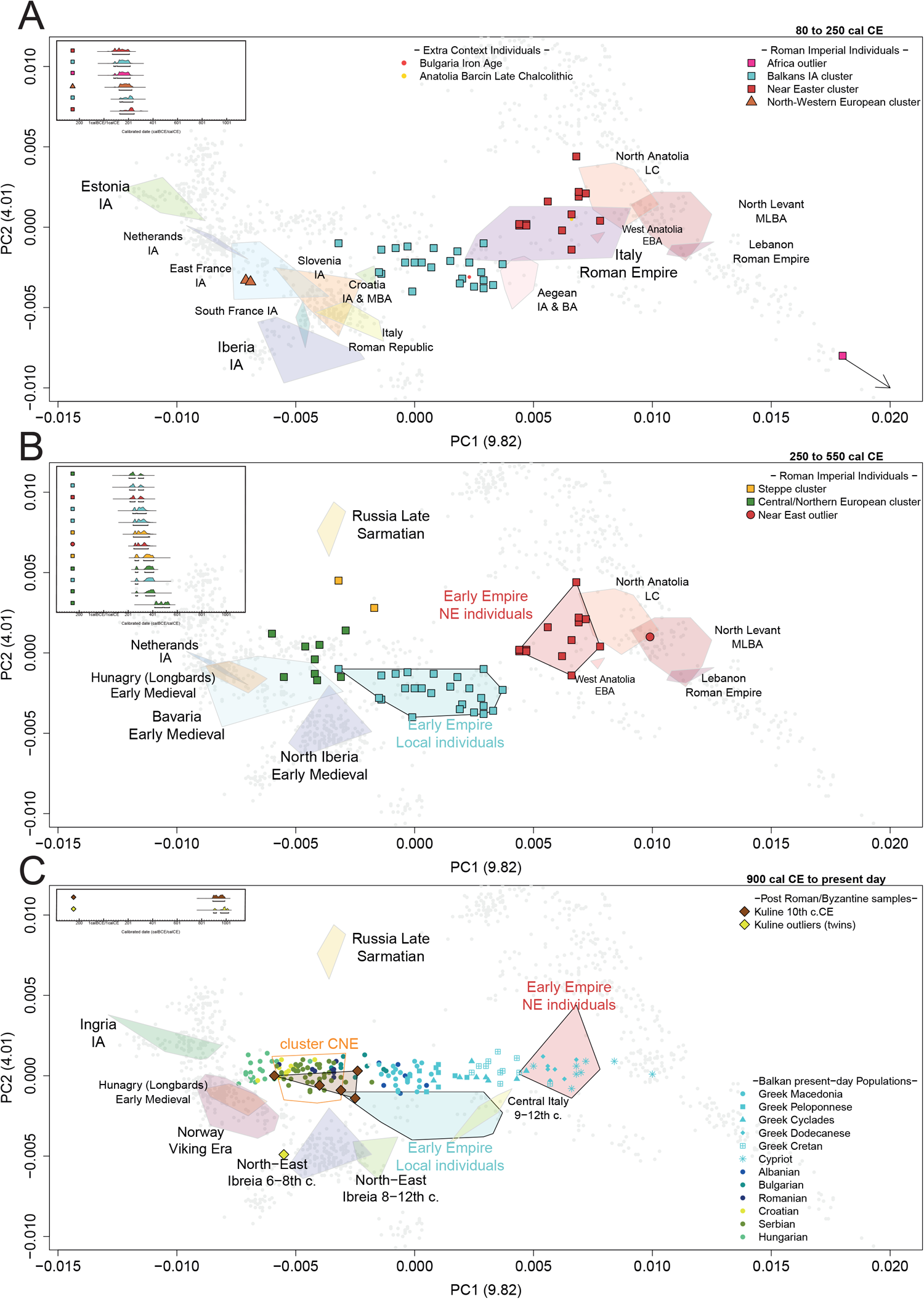
PCA of West Eurasian genetic variability showing present-day individuals as grey circles, published ancient groups as polygons and newly-reported ancient individuals from Viminacium, Timacum Minus and Mediana as colored squares, from A) 0-250 CE, B) 250-500 CE and C) 900 CE to present-day. This PCA is the zoom-in version of Figure S7.

To obtain more insight into the high genetic diversity revealed by PCA and ADMIXTURE (Figure S7 and Figure S10)^14^, we used *qpWave* ^10,15^ to group individuals with similar ancestry composition following a similar approach as in ^16^ (Supplementary section 11). Then, we used *qpAdm* ^10,15^ to model the ancestry of these relatively homogenous groups using previously published earlier, contemporary and later populations from nearby geographical regions (Supplementary section 12). We found 3 major ancestry clusters at Viminacium, Mediana and Timacum Minus Slog and one major ancestry cluster at Timacum Minus Kuline (Figure S11), together with several genetic outliers. In the following sections we place each cluster in its chronological context and provide likely models for its ancestral origin.

### From the founding of the Roman Balkan provinces to the 3^rd^ Century Crisis

At the beginning of the 1^st^ century CE following the conquest of the Dacians, the Roman empire organized its territories in the Balkans as provinces. Viminacium was founded during this century as a military fortification at the Empire’s border and a communications hub along the Danube. The city and region saw a period of prosperity even during the 3^rd^ Century, despite the crisis the Empire underwent during this century due to external pressures and political instability. We found two major clusters of individuals during this period, both present in each of the four necropoli sampled from Viminacium (individuals from Timacum Minus and Mediana date to later periods (Additional Table 1; Table ST1)).

Individuals from the first cluster fall on an area of the PCA delimited by the “ Balkan Iron Age cline” (Figure 1A). Consistent with this, we model the ancestry of this *Balkans Iron Age Cluster* as predominantly deriving from Iron Age (IA) groups from nearby areas in the Balkans, with 67% Aegean Bronze Age-related ancestry and the remainder Slovenia Iron Age-related ancestry (Figure 2; Supplementary section 12.1). A local origin is supported by a high frequency of Y-chromosome lineage E-V13, which has been hypothesized to have experienced a Bronze-to-Iron Age expansion in the Balkans and is found in its highest frequencies in the present-day Balkans ^17^. We interpret this cluster as the descendants of local Balkan Iron Age populations living at Viminacium, where they represented an abundant ancestry group during the Early Imperial and later periods (∼47% of sampled individuals from the 1-550 CE). Excavations of Iron Age Balkans prior to the Roman rule showed the dead where predominantly cremated ^18^, but this changed in Viminacium where inhumation became common suggesting a high degree of Romanization of the local society. Viminacium necropoli followed a bi-ritual mortuary rite where some dead were buried, and some were cremated. During the 1^st^ century until the first half of the 3^rd^ century cremations where more common, however this changed from the 3^rd^ onwards when inhumations prevailed ^19^. We caution that if there was a systematic ancestry difference between the population that buried and the one that burnt its dead, we would of course be obtaining a biased representation of ancestry through ancient DNA analysis.

**Figure 2.**
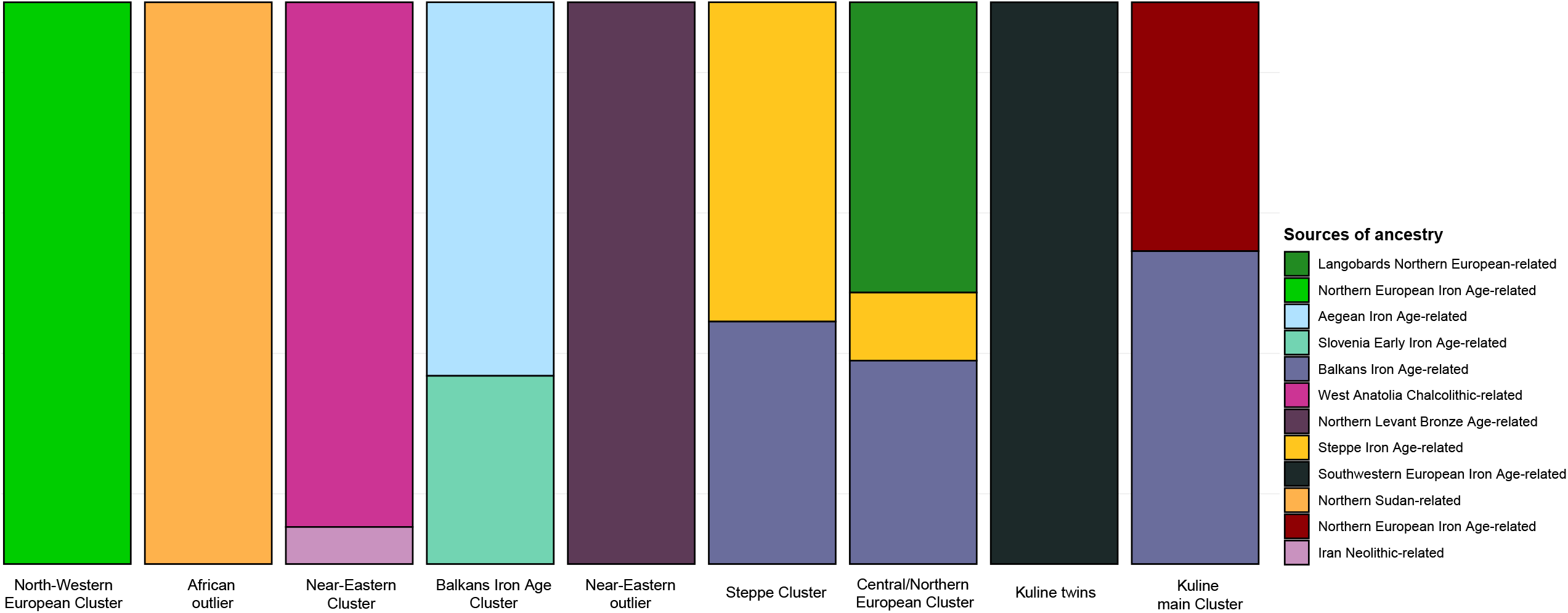
Fitting *qpAdm* models for the different ancestry clusters.

The other major cluster (44% of the samples from Viminacium between 1-250 CE) is represented by individuals who projected towards ancient and present-day Eastern Mediterranean groups in PCA (Figure 1A), close to ancient individuals from Rome during Imperial times ^3^. Their ancestry can be modelled as deriving deeply from Chalcolithic Western Anatolian groups (Figure 2; Supplementary section 12.2), and we refer to this cluster as the *Near Eastern-related cluster*. The same signal of arrivals individuals with Anatolian/Near Eastern ancestral origins is also evident in Rome during the same period ^3^, consistent with large-scale gene-flow originating from the major eastern urban centers of the Empire (such as Constantinople, Antioch, Smyrna and Alexandria). These results suggest that immigration from the east was a common feature across urban centers in the Roman Empire, including in border areas and large cities/military outposts such as Viminacium. Individuals with Eastern Mediterranean ancestry could have high social status: 3 out of the 4 individuals buried in two sarcophagi (each containing a male-female pair) with exceptionally rich grave goods at the Rit necropolis in Viminacium belonged to the *Near Eastern-related* cluster, while the remaining one belonged to the *Balkans Iron Age-related* cluster. This kind of burial was common in the Eastern Roman settlements for aristocratic members of society ^20^. Individuals from this cluster were also more likely to be inhumated in a wooden coffin rather than freely buried, which could also be an indication of higher social prestige.

Three individuals from ∼1-250 CE did not fit into the two major clusters. Two males from Viminacium could be modelled using Iron Age individuals from Northwest Europe as their only source (Figure 2; Supplementary section 12.5), pointing to a Northwestern European origin also supported by the R1b-U106 paternal lineage, which was not been detected in the Balkans in earlier periods but was found at high frequencies in Germanic-speaking areas, both in ancient and present-day individuals. The most remarkable outlier is male I15499, excavated at Pirivoj necropolis in Viminacium, who projects outside West Eurasian genetic diversity (Figure S7). When we incorporated African populations onto the PCA (Figure S8), he projected within the variation of present-day East African populations and close to early Christians from Northern Sudan from 500-800 CE ^21^ who provide a good fit for his ancestry in *qpAdm* (Figure 2; Supplementary section 12.4). An Eastern African ancestral origin agrees with his uniparental markers mtDNA L2a1j and Y-chromosome E1b-V32, both common in East Africa today ^17,22^. Archeological examination of I15499’s grave found an oil lamp depicting an eagle, the symbol of Roman legion (Figure S2C). Although lamps are a common finding in Viminacium graves ^23^, not many depict military iconography. We hypothesize that this male was a Roman legionary or auxiliary stationed at Viminacium. We cannot determine if he was a Roman citizen, although auxiliary military service for a prolonged period of time resulted in citizenship. Historical evidence also points to African recruits being tapped to reinforce the Roman Danubian *limes* ^24^.

### Decline of Roman Rule and the Migration period

During the 3^rd^ century onwards, the Roman Empire was impacted by migrations from the north and east, a process that involved both settlement permitted by the Empire as well as military conflict. It is often viewed as having contributed to the decline of the Western Empire which collapsed politically in 476 CE although Roman control in the Balkans lasted for several more centuries under Byzantine rule. The individuals from the Mediana necropolis, from the Slog necropolis at Timacum Minus, and some individuals at Viminacium coincide with this period (Table ST1). At Slog, we found one directly radiocarbon dated individual with a clear Near Eastern ancestral origin, likely from the Northern Levant (Figure 1B Figure 2; Supplementary section 12.3), as well as directly radiocarbon dated individuals belonging to the *Balkans Iron Age-related* cluster. This confirms that the two major ancestry clusters from 1-250 CE period co-existed at least three centuries in the Danubian *limes*. The legacy of Balkans Iron Age groups persists in admixed form in later groups including present-day Balkan populations (see below), whereas the Near Eastern-related ancestral legacy eventually ebbed in favor of Northern/Eastern European-related ancestry, similar to the patterns observed in the city of Rome itself ^3^. These findings support the hypothesis that such individuals were part of a cosmopolitan group comprising a large proportion of individuals in Imperial towns and cities who over time were demographically overwhelmed by populations in the countryside or by faster reproductive rates of rural or populations without as much Near Eastern influence.

We also observe new ancestry during this period at Mediana, Slog necropolis at Timacum Minus and Viminacium (mostly at Pecine and Vise Grobalja necropoli), as early as the 4^th^ century CE. A cluster of 10 individuals from these necropoli is shifted in PCA from the *Balkans Iron Age-related* cluster toward Central/Northern European ancient and present-day populations (Figure 1B). This group which we refer to as *Central/Northern European cluster*, could be modeled as deriving from two main sources: ∼38% related to the local Balkans Iron Age substratum (we use the *Balkans Iron Age-related* cluster as a proxy for this type of ancestry) and 50% Central/Northern European ancestry (we use as a proxy individuals from a roughly contemporaneous Langobard-associated cemetery in Hungary ^25^). To obtain a fitting model, a significant proportion of ancestry (∼14%) related to contemporaneous nomadic steppe groups (proxied in our analysis by Late Sarmatians from the Eastern Pontic-Caspian steppe ^26^) is also needed (Figure 2; Supplementary section 12.6). This is even more evident in two individuals from the Pecine necropolis in Viminacium (referred to as *Steppe cluster*), who could be modelled as deriving ∼43% of ancestry from the *Balkans Iron Age-related* cluster and 57% ancestry from Late Sarmatian-related Steppe groups (Figure 2; Supplementary section 12.7). Y-chromosome lineages also provide evidence for gene-flow, as 5 of 7 males in the *Central/Northern European* and *Steppe cluster* belonged to two lineages not found in the Balkans earlier: haplogroup I1 with a strong Northern European distribution and haplogroup R1a-Z645, common in the Steppe during the Iron Age and early 1^st^ millennium CE ^26–28^.

The Roman Empire had a prolonged history of contact with Germanic tribes, whose homelands were in Northern Europe between the Rhine and Vistula rivers. During the Great Migration period groups that coalesced as the Goths moved southwards, and settled at the Black Sea north coast prior to their entry in the Roman Empire ^6^. Our observations are consistent with the hypothesis that such tribes interacted with Steppe-related nomadic populations reaching the Eastern European plateau, and incorporated their ancestry into their gene pool before moving into the Balkans. However, the occurrence and manner of this interaction needs to be clarified with a more thorough sampling of this region and time period.

### The formation of the present-day Balkan gene pool

Radiocarbon dating of two individuals from the Kuline necropolis of Timacum Minus places them in the 10^th^ century CE (Table ST1), well after the first Slavic-related settlements in the Balkans during the 7^th^ century ^6^. Two individuals from Kuline were identical twins (Supplementary section 7) and present a different ancestry profile compared to the other individuals from the same necropolis (Figure 1C), likely deriving from Southwestern Europe (Figure 2; Supplementary section 12.9), although they were buried in the same manner as the others from the necropolis ^29^. The remaining five individuals clustered in the West-Eurasian PCA (Figure 1C) on top of the “ present-day Balkan genetic cline”, close to present-day Serbian-speaking individuals that we newly genotyped for this study, but this apparent similarity is a projection artifact as their ancestry could not be fitted using the same *qpAdm* models (Supplementary section 12.8). To understand this, we performed a PCA using present-day Germanic- and Slavic-speaking populations (Supplementary section 9; Figure S9) that we expected would be sensitive to more recent drift separating Central, Northern and Eastern European populations. The Kuline individuals are more shifted towards present-day Slavic-speaking populations as compared to individuals in the *Central/Northern European* cluster, agreeing with the presence of Y-chromosome lineage I2-L621 in Kuline, which is common in present-day Slavic-speaking groups and absent in earlier periods. In light of these results, we modeled the ancestry of the Kuline individuals as a mixture of 56% deriving from the local Balkan Iron Age substratum and 44% deriving from Northeastern European Iron Age groups, and obtained a good statistical fit (Figure 2; Supplementary section 12.8). Our results point to a strong demographic impact of Eastern European groups in the Balkans during the Medieval period, likely associated to the arrival of Slavic-speaking populations. Yet, our results rule out a complete demographic replacement, as we observe a significant portion of local Iron Age Balkan ancestry in Kuline individuals. Interestingly, we found sex bias when modeling the X chromosome of the individuals of this necropolis (Supplementary section 12.8). Perhaps the immigrant groups were constituted by a higher number of women, who therefore impacted more greatly in the demographics of the post-Roman Balkans. However, these findings have only been observed in the Kuline individuals with North-European related ancestry (*n*=5), we suggest more data will be needed to obtain more evidence Slavic sex bias in the Balkans.

To explore whether this Northeastern European ancestry signal persisted in present-day Balkan and Aegean populations, we attempted to model present day groups by using the same *qpAdm* model used for the Kuline individuals (Supplementary section 13). Present-day Serbs, Croats and the rest of central/northern Balkan populations yielded a similar ancestral composition as the Kuline individuals, with approximately 50% Northeastern European-related ancestry admixed with ancestry related to Iron Age native Balkan population (Figure 3), implying substantial population continuity in the region over the last 1,000 years. This ancestry signal significantly decreases in more southern groups, but it is still presents in populations from mainland Greece (∼30%) and even the Aegean islands (7-20%).

**Figure 3.**
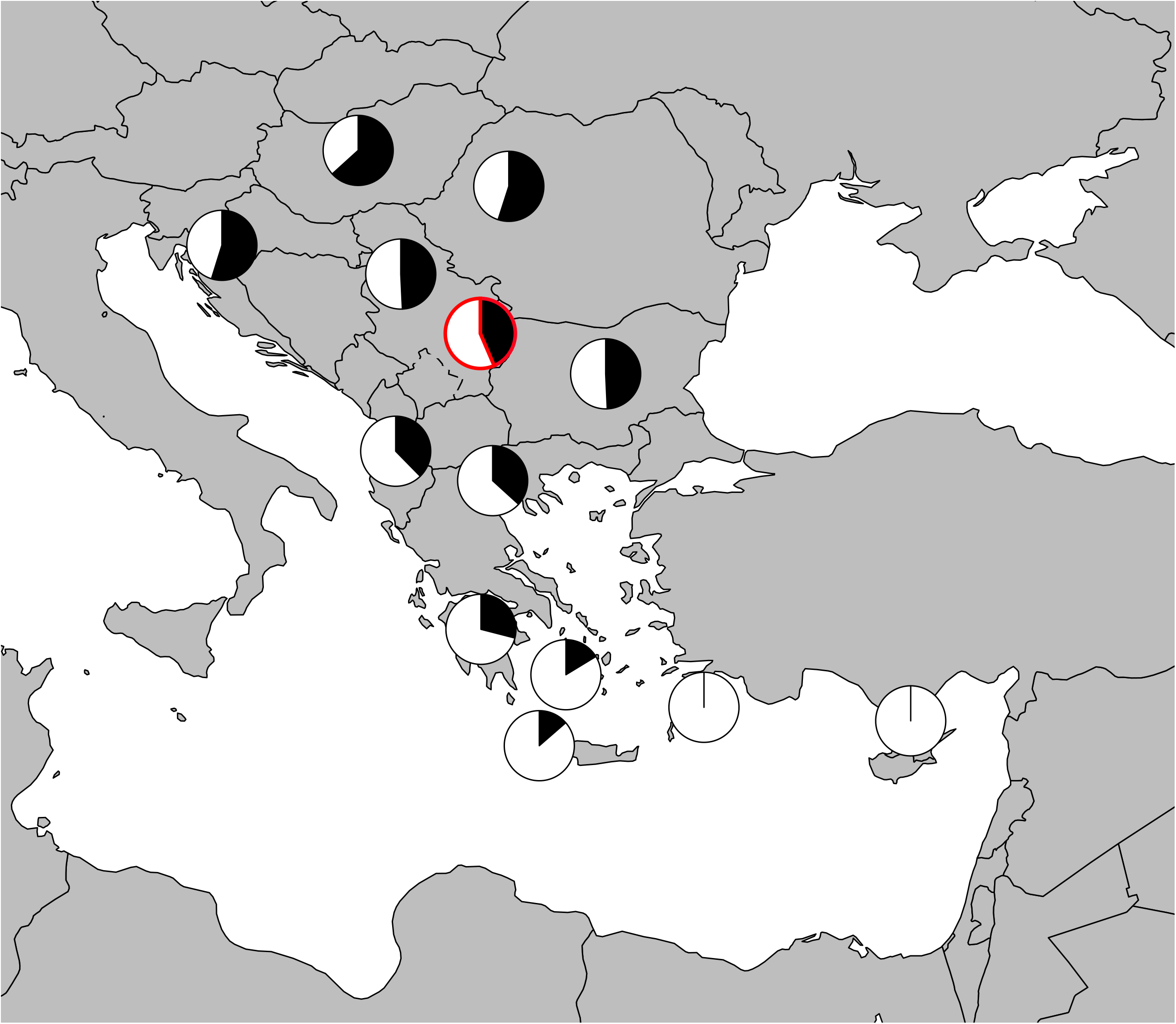
Proportions of Northeastern European-related ancestry (in black) for present-day Balkans populations. 10^th^ century CE Kuline individuals are indicated with a red outline.

## Conclusions

Our results show how people with ancestral origins in Northwestern Europe, Anatolia and East Africa, together with the descendants of Balkan Iron Age populations, inhabited the capital city of the Moesia Province, Viminacium during the Imperial period 1-250 CE (Figure 4A). This indicates a high degree of cosmopolitanism, unprecedented in earlier periods. We found highly similar ancestry trajectories across time in Rome and Viminacium, with a strong Anatolian/Near Eastern influence during the Imperial period that resulted in a large portion of the analyzed individuals in both cities having Near Eastern ancestry, followed by a resurgence of local ancestry after the Empire’s decline ^3^. These results highlight how mobility from the Eastern-most areas of the Empire was a common feature of large cities and towns from the capital city of Rome to the Danubian *limes*, but that demographically these populations were a veneer without long-lasting influences, suggesting either that they were greatly outnumbered by local rural populations, or that their reproductive rates were much lower than that of local rural populations, consistent with evidence that cities and towns in the Roman empire did not successfully reproduce themselves demographically and instead constantly had to be repopulated through immigration ^29^. In the Imperial period, genetic data suggest that a large proportion of this immigration derived from the Eastern Mediterranean highlighting the centrality of this region in the period of intense human connectivity during Imperial Rome. Conversely, the decline in the geographic scale and number of people involved in trans-Mediterranean movements following the Empire’s decline is reflected in the fact that in later periods, Eastern Mediterranean influence largely disappeared in both the city of Rome and in the large towns of the Balkans. An important topic for future ancient DNA research will be to systematically characterize populations in cities and towns as well as rural locations across the Empire, to understand how these patterns differed across geographic locations.

**Figure 4.**
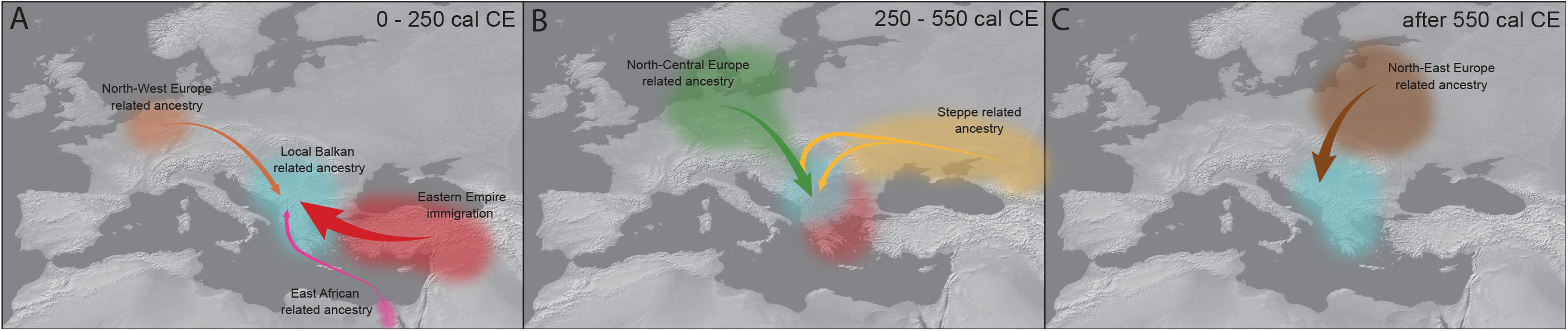
Graphical summary of the genetic findings. A) 0-250 CE, B) 250-500 CE and C) After 550 CE

During and after the fall of the Western Roman Empire, we observe major demographic changes in our sites, with the arrival of ancestry from Northern Europe and the steppe between 250-500 CE, and Northeastern European-related ancestry during the Medieval period, likely associated to the arrival of Slavic-speaking groups (Figure 4C). Our study shows that the Iron Age native Balkan populations were not completely replaced with the arrival of Slavic migrations. These later events generated a cline of populations with relatively similar genetic profiles, yet, three unrelated language groups (Hungarian, Romanian and Southern Slavic) are spoken today in the Central-Northern Balkans area. Our study highlights a decoupling between genetic ancestry and linguistic diversity in modern Balkans that would require further investigation in this critical period. Thorough sampling of Eastern Europe during the first millennium CE will provide a detailed picture of contacts with steppe-nomadic groups as well as the source of the Slavic migrants that made such a key demographic impact on the Balkans.

## Materials and Methods

### Archeological sites

Most of the individuals analyzed (74%) come from the Roman city and military camp of Viminacium, located 12km east of present-day Požarevac and close to the confluence of the Mlava and Danube rivers. It was established in the first half of the 1st century CE, and latter became the biggest settlement and capital of the Upper Moesia (Moesia Superior) province, later First Moesia (Moesia Prima). The importance of the city peaked during the 2^nd^ to the 4^th^ century, with an estimated population of 30,000. The city was sacked by the Huns in 441 and later rebuilt in Justinianic times, but not to its former extent. It was subsequently destroyed by the Avars and abandoned ^11^.

Timacum Minus was a military fort located in the Timok Valley (present-day Serbia). The site is 1km North-East of the village of Ravna, near Beli Timok river. It is believed to be the first fort erected in the 1^st^ century, and was upgraded to a stone fort in the 2^nd^ century, probably during the Trajan era. It is believed that the fort was destroyed in the 441 Hun Balkan campaign ^12^.

During the late Roman Empire, Mediana was a luxurious suburban residence for aristocrats, located next to the town of Naissus. However, this residence was abandoned after Attila’s siege of Naissus in 442 ^13^.

### Ancient DNA Analysis

DNA was extracted from teeth in dedicated clean rooms. The outermost layer of the teeth was removed to reduce possible exogenous DNA contamination; and was drilled at low speed to avoid DNA damage from heat 30. Powder (between 31 and 40 mg per sample) was incubated in lysis buffer and DNA was cleaned and concentrated from one fifth of the lysate following an automated protocol using silica magnetic beads and Dabney Binding Buffer 31. Double-stranded barcoded libraries were prepared with truncated adapters from the extract (corresponding to between 6.2 and 8 mg of powder). All libraries were subjected to partial (‘half’) uracil–DNA–glycosylase (UDG) treatment before blunt-end repair to significantly reduce the characteristic damage pattern of aDNA 32,33.

Two rounds of in-solution enrichment (‘1240k capture’) were performed for a targeted set of 1,237,207 SNPs ^8^ and the complete mitochondrial genome 34. Sequencing was performed by Illumina HiSeq XTen technology for 2 × 101 cycles and 2 × 7 cycles. Mapping was carried out to the hg19 genome assembly using the same tools and process as explained for the mitochondrial capture data.

DNA authenticity was evaluated by estimating the mismatch rate to the consensus mitochondrial sequence, requiring that the rate of damage at the terminal nucleotide was at least 3% for partial UDG-treated libraries, examining the ratio of X-to-Y chromosome reads, and estimating X-chromosome contamination in males based on the rate of heterozygosity.

### Mitochondrial haplogroup determination

Reads mapped to mitochondrial reference genome were used to determine mtDNA haplogroups (using sequences with mapping quality (MAPQ) ≥ 30 and base quality ≥ 30). A consensus sequence was first using bcftools and SAMTools 35 using a majority rule and requiring a minimum coverage of two 36. These consensus sequences were then used to call mitochondrial haplogroups using *HaploGrep2* based on phylotree (mtDNA tree build 17) 37,38.

### Y-chromosome analysis

Using sequences with MAPQ ≥ 30 and a base quality ≥ 30 36 mapping to 1240k Y-chromosome targets, haplogroups were called by determining the most derived mutation for each sample using the nomenclature of the International Society of Genetic Genealogy (http://www.isogg.org; version 15.73) 36.

### Sample sex determination and kinship analysis

Sample sex was determined by computing the ratio of Y-chromosome reads to the sum of chromosomes X and Y reads. We used READ (Relationship Estimation from Ancient DNA) program 39 to determine sample kinships.

### Merge of newly generated data with published data

Two datasets were assembled for genome-wide analyses (Supplementary section 8). We created these datasets by merging the newly reported ancient and modern individuals with published present-day and ancient individuals.

The first dataset includes 2562 present-day individuals from worldwide populations which were genotyped on the Human Origins Array 10,40–42.

The second dataset includes the same ancient samples but merged with 300 present-day individuals from 142 populations sequenced by the Simons Genome Diversity Project to a high coverage 43.

### Principal Component Analysis

Principal Component analyses (PCAs) were performed using the *smartpca* module of the EIGENSOFT software (Version 7.2.1). We computed principal components on 1027 present-day West Eurasian individuals for the West-Eurasia (WE) PCA; on 1226 present-day west Eurasians, North-Africans and Sub-Saharan Africans for the WE+Africa PCA; and on 407 present-day individuals for the Northern-Europe PCA. Ancient individuals were projected onto the components using lsqproject:YES and shrinkmode:YES 44,45 (https://www.hsph.harvard.edu/alkes-price/software/).

### ADMIXTURE analysis

ADMIXTURE (version 1.3.0) ^14^ was used to compute proportions of putative ancestral components (*K*) present in each one of the individuals. ADMIXTURE analysis was performed on the ‘HO’ dataset (Supplementary section 8).

### *f*-statistics

*f*-statistics were computed using ADMIXTOOLS ^10^ with default parameters. We used *qpWave* with the option allsnps:YES. This tool was used to group individuals into clusters of shared ancestry, which can be treated as a coherent group in *qpAdm* (Supplementary Section 11). The outgroup population list used for this test can be found in Table ST2.

We used *qpAdm* test to model the ancestry of each of the sample-clusters, with allsnps:YES. The list of populations used for these computations can be found in (Additional Table 3). We computed standard errors for both *qpWave* and *qpAdm* using a weighted block jackknife over 5-Mb blocks (Supplentary section 12).

## Author Contributions

I.O., P.C., C.L-F., M.G, and D.R conceived and designed the study. I.O. and P.C. performed the genetic data analysis and wrote the paper with the help of all authors. C.L-F., D.R., I.M. and M.G. supervised the study, and contributed to the writing of the paper. I.M., M.K., S.G., S.P., N. M-R. and D. V. performed the archeological excavations and sample extraction and provided the archeological and anthropological context. N.R, A.C., N.A., K.S, A.M.L, F.Z. and K.C performed the laboratory work. S.M performed bioinformatics data processing. I.L, performed analysis on the uniparental markers and contributed to the design of analytical steps. Z.T, D.K. and M.G. provided and sampled genetic information of the newly reported present-day individuals and contributed to the result interpretation.

## Acknowledgments

We thank V. Villalba-Mouco for advice and comments on the manuscript, A. Claxton and N. Adamski for laboratory work, and M. Coronado-Zamora for assistance in Figure 4. This research was partially supported by a PGC2018-0955931-B-100 grant (MCIU/AEI/FEDER, UE) of the Spanish Ministry of Science of Innovation, awarded to C.L.-F, and by a fellowship from “ la Caixa” Foundation (ID 100010434), code LCF/BQ/PI19/11690004, awarded to I.O. P.C. is supported by an FPI-2019 (BDNS ID:476421) grant of the Spanish Ministry of Science of Innovation. I.O. is supported by a Ramón y Cajal grant from Ministerio de Ciencia e Innovación, Spanish Government (RYC2019-027909-I). D.R. was supported by the National Institutes of Health (NIGMS GM100233), the John Templeton Foundation (grant 61220), and by the Allen Discovery Center program, a Paul G. Allen Frontiers Group advised program of the Paul G. Allen Family Foundation; D.R. is also an Investigator of the Howard Hughes Medical Institute.

