## Supplementary Material for "Cosmopolitanism at the Roman Danubian Frontier, Slavic Migrations, and the Genomic Formation of Modern Balkan Peoples"

**Contents**

1. **Present-day newly reported individuals**
2. **Archeological context of newly reported individuals**
3. **Direct AMS ^14^C Bone Dates**
4. **Ancient DNA laboratory work**
5. **Bioinformatics processing**
6. **Mitochondrial and Y- chromosome haplogroup determination**
7. **Kinship analysis, sex determination, aneuploidies and ROHs**
8. **Genome-wide analysis datasets**
9. **Principal Component Analysis**
10. **ADMIXTURE**
11. ***qpWave* admixture testing**
12. ***qpAdm* admixture modeling**
13. ***qpAdm* admixture modeling of present-day Balkan populations**
1. **Present-day newly reported individuals**

We collected genetic material from 37 unrelated Serb individuals from Serbia (*n=*19), Montenegro (*n*=7), Croatia (*n=*5), North Macedonia (*n*=1) and Bosnia and Herzegovina (*n*=5). The sample collection and genotyping were carried out with the approval and accordance to the supervision of the Ethical committee of the Institute for Molecular Genetics and Genetic Engineering, University of Belgrade (O-EO-29/2021). Participants were informed about the goals of the project and gave informed consent. Genomic information was obtained by Affymetrix Human Origins Array genotyping of DNA extracted from buccal tissue ^1^, with data quality control performed as described previously ^2^. Additional File Table 2 shows information from these individuals.

2. **Archeological context of newly reported individuals**

**2.1 Viminacium**

Viminacium was the capital of Upper Moesia, a Roman province in the territory of present-day Serbia. The military significance of Viminacium lay in its strategically position on the Danube River, as a military camp housing two legions: *Legio VII Claudia pia Fidelis* and *Legio IV Flavia*. This city saw a period of prosperity during the 2^nd^ century and developed into the biggest urban settlement in Upper Moesia. This progress was in part due to Viminacium’s strategic location at the mouth of the Mlava River, who acted as port for the exportation of Mlava Valley’s agricultural and mining goods across the Danube. Under Emperor Gordian III's rule, the city received the status of colony, with the right to mint coins. The city also occupied an important military strategic location as it was an important connection hub. Viminacium acted as a crossroads which connected the north of the Balkan peninsula with the roman controlled West and South. The Pannonia Road also passed through the city, which crossed the Danube and ended in the Black Sea ^3^.

Several necropolises from the Viminaium archaeological site have been found and studied. Physical anthropological from the recovered individuals at analysis at Viminacium’s necropolises has pointed to diverse morphological profiles ^4–6^. The orientation of the graves varied from the direction E-W or W-E to N-S or S-N with deviations. In some cases, grave orientation could be related to religious beliefs, such as pagan or Christian beliefs which coexisted in Viminacium. A map of Viminacium archeological excavations can be found in **Figure S1A**, together with its location in the Balkans **Figure S1B.**

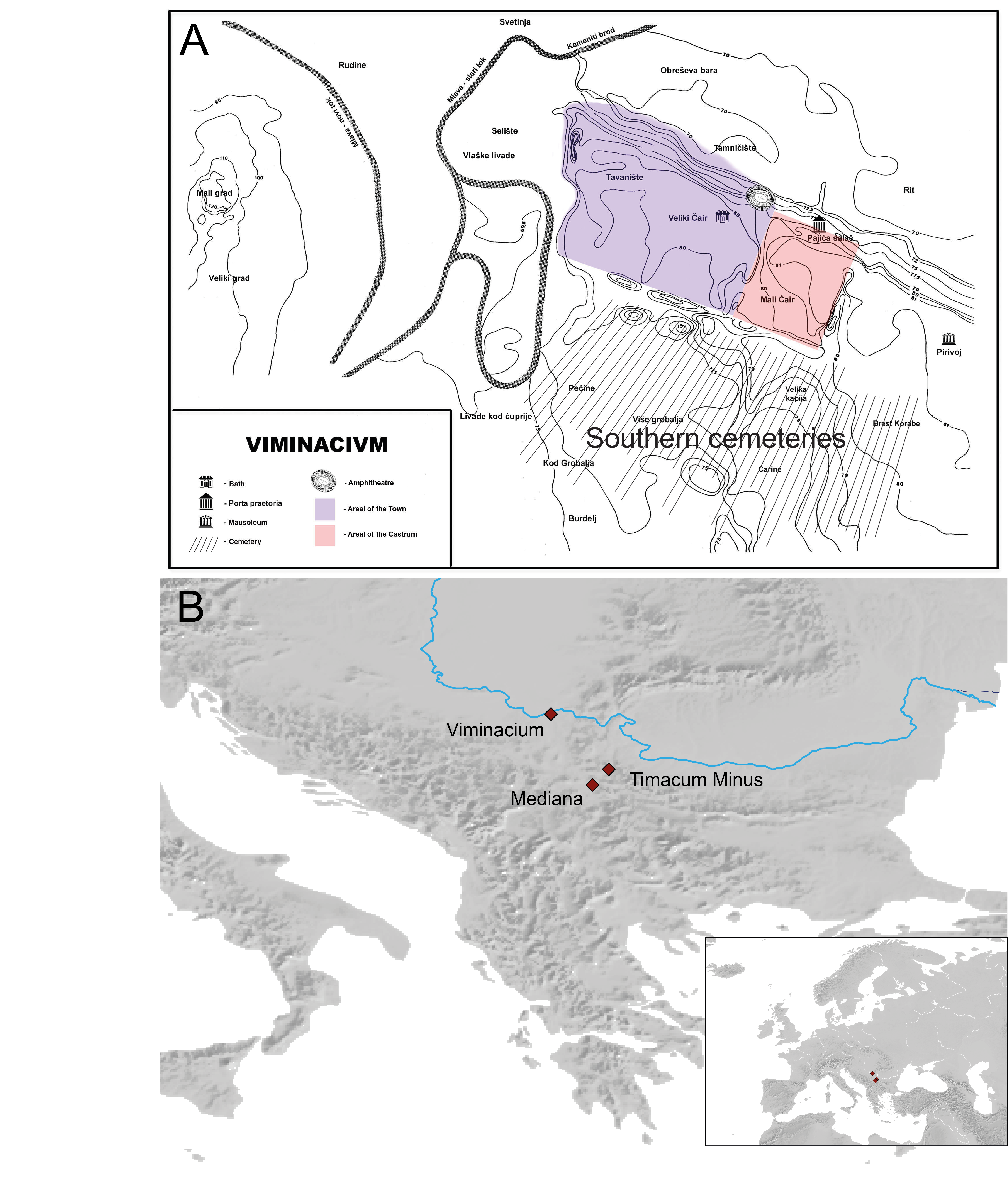

**Figure S1. A)** Viminacium archeological site. **B)** Location of the 3 archeological sites from this study.

In what follows, we describe the archaeological context and inventory of the graves selected for ancient DNA analysis. Individuals were chosen from four necropolises based on archaeological context, dating and skeletal preservation.

**2.1.1 Pećine necropolis**:

The necropolis of Pećine is the largest on Viminacium and archaeological research on it is still ongoing. So far, over 6,000 graves from the prehistoric periods (including a Celtic cemetery) to the Middle Ages have been found. For this project we selected samples from cemeteries were archeologically dated to the 1st to 4th century CE. All the deceased can be classified as having had a modest type of burial, so they are attributed to the rather poorer citizens of Viminacium. Archaeological finds are also modest. Graves 5757 and 5762 (not sampled for ancient DNA) can be classified according to the finding of coins in the period of the fourth decade of the 4th century CE, and according to the west-east and east-west orientation they might be early Christians. This agrees well with two radiocarbon dates from G-5665 (258-413 cal CE (1700±20 BP, PSUAMS-8554)); and G-5736 (246-365 cal CE (1745±15 BP, PSUAMS-8591)). Another radiocarbon date from G-2771 (70-208 cal CE (1910±20 BP, PSUAMS-8553)) yielded an earlier date within the 2^nd^ century CE, implying a long-term use of the necropolis (**Table ST1**; **Additional Table 1**). Burial practices are typical of the Roman period at Viminacium (grave with no construction, deceased in wooden coffin or graves with brick construction with the seal of the legion).

Sampled Graves:

Freely buried deceased: G-5665 (S-N), G-5703 (N-S), G-5736 (E-W), G-2771 (W-E), G-4661 (Burial in prehistoric pit, S-N), G-5924 (NW-SE), G-5769 (NW-SE).

Deceased in a wooden coffin: G-5752 (E-W), G-3082 (SE-NW), G-3053 (NE-SW).

**2.1.2 Pirivoj necropolis:**

The Pirivoj necropolis, among other features, contains the presumed mausoleum of Emperor Hostilian, who was likely cremated there. About 450 inhumed skeletons have been excavated and anthropologically analyzed. In addition to the necropolis, the remains of non-religious buildings have been excavated. According to the archaeological findings, people buried there and sampled for this project are citizens of a more modest and middle class who can be dated to the period from the 2nd to 4th century, confirmed with radiocarbon dates of G-103 (80-215 cal CE (1895±20 BP, PSUAMS-8552)) and G-402 (124-228 cal CE (1870±20 BP, PSUAMS-8590)) (**Table ST1**; **Additional Table 1**). We sampled grave G-103 whose deceased was placed in a wooden coffin. The tomb contained an oil-lamp with an imperial eagle, a traditional Roman military symbol (**Figure S2C**). The individual recovered from this grave had African ancestral origins (see Supplementary section 12.4).

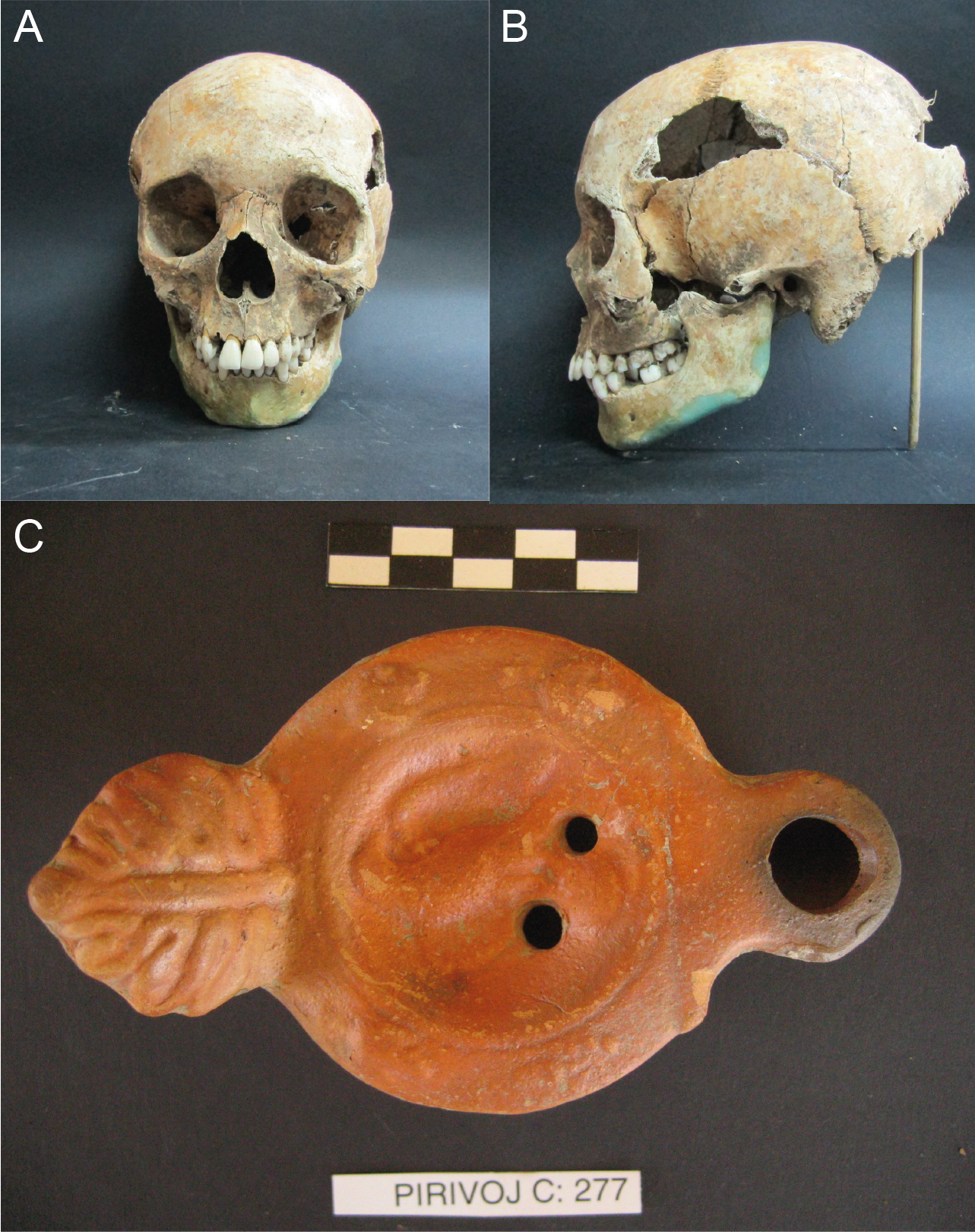

**Figure S2**. Elements recovered from Grave G-103 in Pirivoj Necropolis (Viminacium). **A)** Frontal view of I15499 skull. **B)** Lateral view of I15499 skull **C)** Oil lamp with a Roman Eagle.

Sampled Graves:

Square construction tomb: G-12

Freely buried deceased: G-61 (W-E), G-322 (S-N), G-314 (W-E), G-341 (S-N), G-357 (NE-SW).

Deceased in a wooden coffin: G-101 (W-E), G-104(NW-SE), G-103, G-105, (E-W), G-353 (W-E), G-361 (N-S), G-402 (NW-SE)

Deceased in graves with brick construction: G-36 (W-E), G-47 (W-E), G-288 (N-S).

**2.1.3 Rit necropolis:**

The Rit necropolis was excavated as part of a settlement that contained Roman villas and other non-religious buildings. A total of 150 inhumed skeletons were examined. In addition to the burials that are characteristic of the Roman population on Viminacium, this necropolis also features burials with a clear Eastern influence such as two sarcophagi (G30 and G148) (**Figure S3**) each containing a male-female pair and rich grave goods that can be dated to the period from the 1st to the 3rd century CE. Three out of four individuals found in these two sarcophagi yielded Near Eastern ancestry (further explained in Supplementary section 12.2).

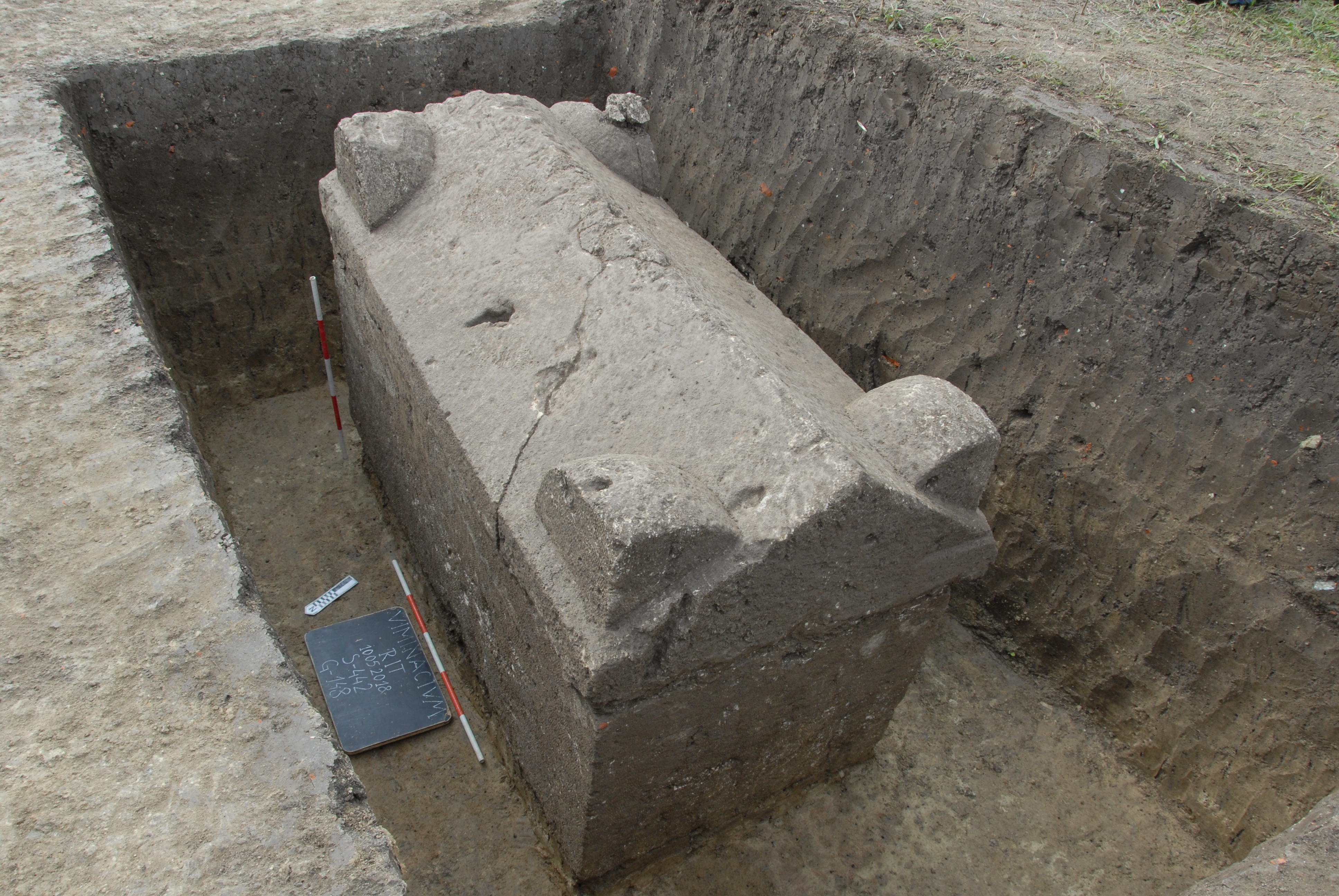

**Figure S3.** Sarcophagus of Grave 148 at the Rit Necropolis (Viminacium).

According to the findings of coins (Graves G-147 and G-112) with the image of Antoninus Pius issued during his rule in the 2^nd^ century, which means that they could represent some of the oldest inhabitants of Viminacium. The Radiocarbon dating of G-103 from this cementery (129-247 cal CE (1835±20 BP, PSUAMS-8560)) supports the chronological attribution.

Sampled Graves:

Freely buried deceased: G-81 (W-E), G-112 (S-N), G-127 (W-E)

Deceased in a wooden coffin: G- 23 (W-E), G-147 (W-E)

Deceased in graves with brick construction: G-11 (W-E), G-84 (W-E), G-122, G-65 (W-E), G-123

Sarcophagi: G-30 (W-E), G-148 (W-E).

**2.1.4 Više Grobalja necropolis:**

The necropolis of Više Grobalja is the second largest in Viminacium, where archeological excavations are still ongoing and about 2,850 inhumed graves have been excavated so far. As in the case with Pećine necropolis, archaeological excavations are of protective character for the carbon needed to feed Drmno thermal power plant. Individuals from this necropolis belong to a somewhat average group that practiced the burial method characteristic of most of the deceased from the Roman period in Viminacium.

Sampled Graves:

Freely buried deceased: G-105 (NE-SW), G-597 (W-E), G-916 (N-S), G-1614 (NW-SE), G-962.

Deceased in a wooden coffin: G-2307 (W-E), G-343 (N-S), G-2012 (W-E), G-2292 (E-W), G-351 (NE-SW)

Deceased in graves with brick construction: G-1428 (W-E) seal LEGVIICL.

**2.2 Timacum Minus**

Timacum Minus was founded as a military fortification defending the left bank of the Beli Timok River. A small village developed adjacent to the fort. The site is located in the vicinity of present-day Ravna, 10 km north of Knjaževac (Serbia). There are several hypotheses regarding the nature of this Roman site, however, the most credible assumption is that it was a fortified administrative center of the Upper Moesia mining region *Territoria metallorum*, which comprised the north-eastern part of the Upper Moesia province, later the provinces of Dacia Ripensis and Dacia Mediterranea ^7^.

Two necropolises were sampled at the Timacum Minus Archeological site. At both necropolises the archaeological findings suggests they contained military graves as they had a significant amount of weapons.

**2.2.1 Slog Necropolis**

This necropolis was formed in three successive phases, from the middle of the 4th to the middle of the 5th century: phase I, from;, from 350 to 380 CE ,phase II– from 380 to 410 CE– and phase III, from 410 to 450 CE ^7^. When it comes to the Roman period and Early Medieval, the following samples were taken from the necropolis Slog according to their proposed necropolis phase:

Sampled Individuals:

From PHASE I of the Necropolis (350–380 CE):

G-91 A destroyed grave construction of stone and bricks (W-E)

G-99 Rectangular burial pit dug into the subsoil (W-E).

From PHASE II of the Necropolis (380–410 CE):

G-15 Rectangular burial pit dug into the subsoil, above which there was a construction of broken stone and tegulae that was destroyed and dislocated next to the grave.

G-25 Grave construction made of river pebbles, preserved on both sides of the head, below the feet and along the right side of the skeleton.

G-26 Destroyed grave construction of river pebbles above the rectangular burial pit. Finds of nails, (below the legs and along the left side of the skeleton, indicate the existence of a wooden casket.

G-27A Destroyed rectangular construction of river pebbles above a burial pit.

G-28 Destroyed grave construction of river pebbles above a burial pit.

G-97 Remains of a stone construction around the skull (W-E).

G-123 Freely dug burial pit (W-E).

From PHASE III of the Necropolis (410–450 CE):

G-100 Destroyed grave construction of stone and brick joined with mortar (W-E)

G-108 Rectangular grave construction, a cist of tegulae laid edgewise.

**2.2.2 Kuline Necropolis**

Archeological excavations in Ravna Village have found 4 different medieval necropoli: 1- Kuline, located in the interior of the Roman fortress, 2-Slog, 3- Podina and 4- Ravanski (Zubanov).

The excavations of Kuline necropolis started in 1978. A total of nine graves were initially found, located in the central part of the Roman fortress. The necropolis is organized in rows, which are oriented in the west-east direction, and are formed by grave pits, except in two cases where river pebbles were found along the left side of the deceased ^8^. A total of 7 individuals were sampled:

G-1 Freely burial deceased (W-E)

G-2 A simple grave pit was in the cobblestones of the street (W-E).

G-3 As in previous grave, the pit was in the cobblestones of the street (W-E)

G-4 Freely buried deceased (W-E).

G-5 Freely buried deceased (W-E).

G-6 Freely buried deceased (W-E).

G-7 freely buried deceased (W-E).

**2.3 Mediana**

Mediana is a Roman archaeological site located on the left bank of the river Nišava and on the old road Niš-Pirot near the village of Brzi Brod (Serbia). The remains are located approximately 4,5 km east of the Niš Fortress (below which the ancient city of Naissus can be found). During antiquity it was a suburb that served to supply the ancient Naissus. Emperor Constantine the Great, who was born in Naissus, remodeled Mediana to construct a luxurious residential villa. The site was located on the old Roman *via publica* that passed through the diocese Dacia in direction *Singidunum – Viminacium – Naissus – Serdica* ^9^.

Archeological excavations during the years 2000-2001 unearthed 4 graves (Graves 34 to 37) ^10^. Graves 34 (located in probe 4) and 35 (pro-expansion of grave 34) were found at the west of the villa, in the peristyle. The archeological findings in grave 34 (a rowcomb, a lunular bronze pendant and a bead of glass paste), indicates it can be dated to the end of the 4th - beginning of the 5th century. This tomb was constructed re-using Roman bricks. Grave 35, on the other hand, was carved into a stone column from the end of the 4^th^ century – beginning of the 5^th^ century. This column therefore belongs to the latest construction phase of the villa and is oriented quite differently in relation to other architectural units. This tomb contained a bronze coin (held in the right hand), an iron thread and an iron knife.

Both skeletons laid in their graves in an outstretched position, with their hands alongside their bodies. They had an east-west orientation, with their heads on the west ^4^. The archaeological and historical context indicates these individuals belonged to the Gothic cultural circle ^9^. In connection with this, both individuals present artificially deformed skulls which were characteristic of Germanic tribes during the Great Migration period (**Figure S4**). This cultural practice was could have been adopted from the Huns ^11^. Furthermore, the skull bandaging method observed for these two individuals corresponds with other examples of cranial deformation previously observed in Germanic necropolises at Viminacium ^12^. There are 4 bandaging zones observed form a lateral projection: frontal above the measuring point glabella (G), parietal behind the bregma (B), parietal above the inion (I), as well as occipital below the measuring point inion. Thus, in a reconstructive sense, it is obvious that the bandage tape bent around the cerebral part of the skull in three directions: fronto-parietal, fronto-occipital and parieto-occipital ^4^.

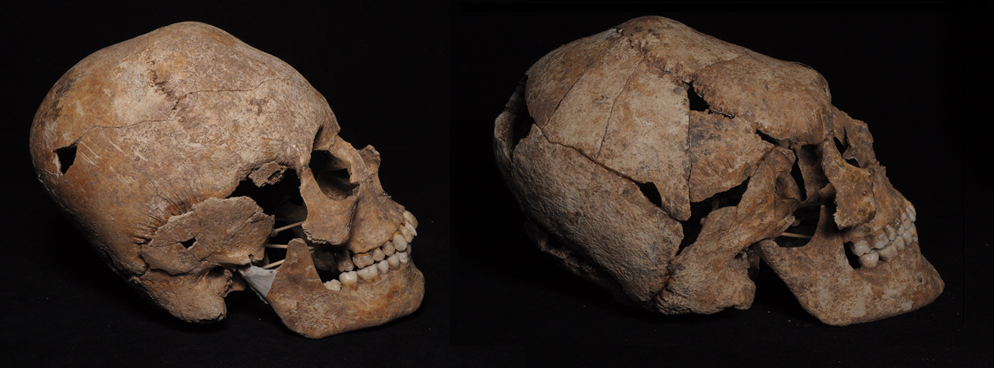

**Figure S4.** Elongated skulls of Graves 34 (left) and 35 (right), unearthed at the Mediana archeological site.

From this necropolis, only G-34 individual yielded enough DNA for further genetic study. The analysis of this sample points towards them having Central-North European Iron Age- related ancestry, admixed with a small yet significant portion of Iron Age Steppe related ancestry, compatible with possible Germanic peoples (further explained in Supplementary section 12.6).

Sampled Graves:

G-34

G-35

3. **Direct AMS ^14^C Bone Dates**

Radiocarbon dating was performed at the Pennsylvania State accelerator mass spectrometry radiocarbon laboratory as described in previous publications ^13,14^.

**Table ST1.** Radiocarbon dating results on the newly reported samples carried out specifically for this study. Genetic cluster attribution is further explained in supplementary section 12.

| **Master ID** | **Date** | **Genetic cluster** | **Necropolis** |
| --- | --- | --- | --- |
| I15527 | 70-208 calCE (1910±20 BP, PSUAMS-8553) | North-Western European cluster | Viminacium, Pecine Necropolis |
| I15499 | 80-215 calCE (1895±20 BP, PSUAMS-8552) | African outlier | Viminacium, Pirivoj Necropolis |
| I15528 | 80-215 calCE (1895±20 BP, PSUAMS-9562) | Near Eastern  cluster | Viminacium, Pecine Necropolis |
| I15517 | 124-228 calCE (1870±20 BP, PSUAMS-8590) | Near Eastern  cluster | Viminacium, Pirivoj Necropolis |
| I15521 | 130-237 calCE (1850±15 BP, PSUAMS-9560) | Balkans IA cluster | Viminacium, Vise Grobalja  Necropolis |
| I15500 | 129-247 calCE (1835±20 BP, PSUAMS-8560) | Balkans IA cluster | Viminacium, Rit Necropolis |
| I15520 | 207-326 calCE (1810±20 BP, PSUAMS-9559) | Central/Northern Europe cluster | Viminacium, Vise Grobalja  Necropolis |
| I15518 | 211-326 calCE (1805±20 BP, PSUAMS-9558) | Balkans IA cluster | Viminacium, Rit Necropolis |
| I15526 | 215-326 calCE (1800±20 BP, PSUAMS-9561) | Near Eastern  cluster | Viminacium, Vise Grobalja  Necropolis |
| I15534 | 237-348 calCE (1765±20 BP, PSUAMS-9563) | Balkans IA cluster | Viminacium, Pecine Necropolis |
| I15492 | 241-362 calCE (1755±20 BP, PSUAMS-9557) | Balkans IA cluster | Viminacium, Pirivoj Necropolis |
| I15551 | 242-375 calCE (1750±20 BP, PSUAMS-8561) | Near Eastern outlier | Timacum Minus, Slog Necropolis |
| I15533 | 246-365 calCE (1745±15 BP, PSUAMS-8591) | Asian ancestry | Viminacium, Pecine Necropolis |
| I15535 | 255-409 calCE (1715±20 BP, PSUAMS-9564) | Asian ancestry | Viminacium, Pecine Necropolis |
| I15549 | 259-409 calCE (1705±15 BP, PSUAMS-8557) | Central/Northern Europe cluster | Mediana |
| I15531 | 258-413 calCE (1700±20 BP, PSUAMS-8554) | Central/Northern Europe cluster | Viminacium, Pecine Necropolis |
| I15544 | 261-418 calCE (1685±20 BP, PSUAMS-8725) | Balkans IA cluster | Timacum Minus, Slog Necropolis |
| I15545 | 417-538 calCE (1610±15 BP, PSUAMS-8556) | Central/Northern Europe cluster | Timacum Minus, Slog Necropolis |
| I15538 | 892-989 calCE (1115±15 BP, PSUAMS-8592) | Kuline outlier (Twins) | Timacum Minus, Kuline  Necropolis |
| I15542 | 897-1021 calCE (1075±15 BP, PSUAMS-8555) | Kuline 10th Cen. | Timacum Minus, Kuline  Necropolis |

4. **Ancient DNA laboratory work**

We performed laboratory work in dedicated clean rooms at the Reich lab (Harvard Medical School). The outermost layer of the teeth was removed to reduce possible exogenous DNA contamination; and was drilled at low speed to avoid DNA damage from heat ^15^.

We extracted DNA, using silica magnetic beads and Dabney binding buffer ^16^ and prepared double-stranded barcoded libraries with truncated adapters. Libraries were subjected to partial (‘half’) uracil–DNA–glycosylase (UDG) treatment before blunt-end repair to significantly reduce the characteristic damage pattern of aDNA ^17,18^.

DNA libraries were enriched for human DNA using probes that target 1,233,013 SNPs (‘1240k capture’) ^19,20^ and the mitochondrial genome. Captured libraries were sequenced on an Illumina HiSeq Xten instrument with 2x101 cycles and 2x7 cycles to read out the two indices ^21^.

5. **Bioinformatics processing**

Reads for each sample were extracted from raw sequence data according to sample-specific indices added during wetlab processing, allowing for one mismatch. Adapters were trimmed and paired-end sequences were merged into single ended sequences requiring 15 base pair overlap (allowing one mismatch) using a modified version of *SeqPrep 1.1* (https://github.com/jstjohn/SeqPrep) which selects the highest quality base in the merged region. Unmerged reads are discarded prior to alignment to both the human reference genome (hg19) and the RSRS version of the mitochondrial genome using the ‘samse’ command in *bwa* (version 0.6.1) ^22^. Duplicates were removed based on the alignment coordinates of aligned reads, as well as their orientation. Libraries were sequenced to saturation across multiple sequencing lanes where necessary, with complexity metrics established using *preseq* ^23^, merging where necessary. Subsequent authenticity of ancient DNA is established using several criteria: we discarded from further analysis libraries with a rate of deamination at the terminal nucleotide below 3%. We computed the ratio of X-to-Y chromosome reads, estimated mismatch rates to the consensus mitochondrial sequence, using *contamMix-1.0.10* ^24^ and ran X-chromosome contamination estimates using *ANGSD* ^25^ in males with sufficient coverage (Table S2). Libraries with evidence of contamination were discarded from genome-wide analyses or, in cases with sufficient data, restricted to sequences with cytosine deamination to remove potential contaminating sequences. We required a minimum of 20,000 SNPs with at least one overlapping sequence for inclusion in genome-wide analyses.

6. **Mitochondrial and Y- chromosome haplogroup determination**

To determine mtDNA haplogroups from the ancient samples, reads were mapped to mitochondrial reference genome. Sequences with mapping quality (MAPQ) ≥ 30 and base quality ≥ 30 were used for the determination. A consensus sequence was first constructed using *bcftools* and *SAMTools* ^22^ using a majority rule and requiring a minimum coverage of two ^26^. These consensus sequences were then used to call mitochondrial haplogroups using HaploGrep2 based on phylotree (mtDNA tree build 17) ^27,28^.

Similarly, we used sequences with MAPQ ≥ 30 and a base quality ≥ 30 to determine the Y chromosome lineages from the male ancient individuals:

- We annotated the haplogroup associated with the most derived mutation for each sample using the nomenclature of the International Society of Genetic Genealogy (http://www.isogg.org; version 14.76; Accessed April 25^th^ 2019).

- We annotated the path of derived mutations for each sample using the nomenclature of Yfull 8.09 (<https://www.yfull.com/>) as described in Lazaridis et al., in submission.

The results of the mtDNA and Y-chromosome haplogroup determinations can be found in Additional file Table 1.

7. **Kinship analysis, sex determination, aneuploidies and ROH**

We tested for kinship relationships among pairs of individuals included in our study. To do this, we used the Relationship Estimation from Ancient DNA (READ) program implemented by ^29^, which can infer family relationships up to second degree even from samples with very low coverage. Degrees of kinship classification in a population must be independent of within0population diversity, and thus the proportion of non-matching alleles (P0) needs to be normalized before classifying relationships between pairs of individuals. This is achieved by using the expected value for a randomly chosen pair of unrelated individuals from the same population. Both window size and median pairwise P0 default options were used. We found two close kinship relationships. The first is between two individuals (I15490 and I15491) from Pirivoj Necropolis (Viminacium) buried in the same double grave G-36 who are likely second-degree relatives. Given that they have different mtDNA lineages and the same the Y-chromosome lineage, they could be paternal half-brothers, nephew and paternal uncle, or grandson and grandfather. The other kinship relationship was between I15538 (Grave 2) and I15539 (Grave 3) from Kuline Necropolis at Timacum Minus who were found to be identical twins. In order to validate this result and disregard any possible mix-up during sampling or laboratory analysis, we took a new sample from each individual targeting the same dental piece in both cases to minimize the possibility of sampling two teeth from the same skeleton. The new sample from Grave 3 (I26846) was again determined to be a genetic duplicate of I15538 and I15539 using READ, while the new sample from Grave 2 (I26846) failed genome-wide 1240k capture analysis but the mtDNA belongs to the same rare mtDNA haplogroup H1e1a6 as the other three samples. Altogether, this evidence strongly suggests that the individuals from Graves 2 and 3 are indeed identical twins (**Figure S5)**.

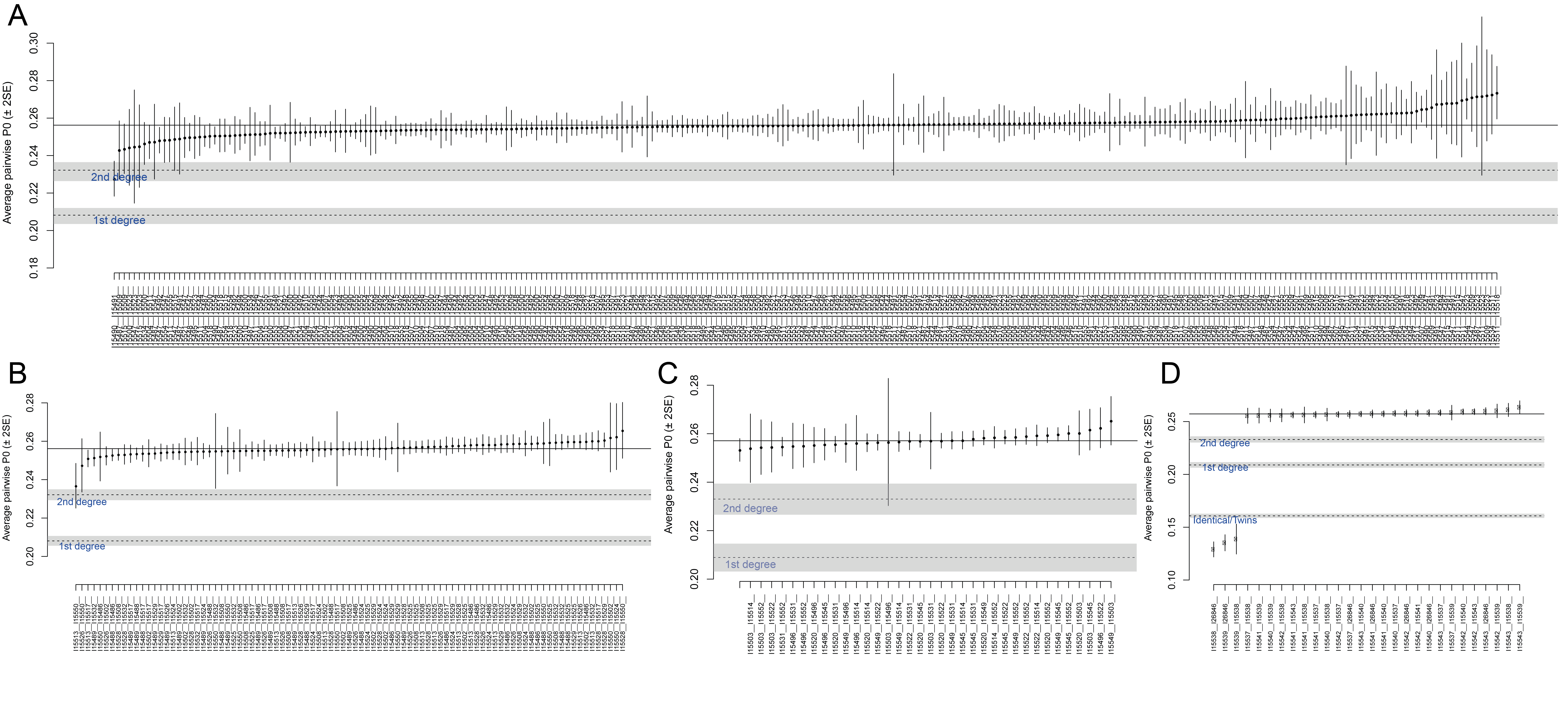

**Figure S5.** Kinship relationships exploration among the newly reported ancient samples, divided by shared ancestry clusters. A) Comparisons within the local Balkan Iron Age cluster. B) Comparisons within the Central Northern European ancestry cluster. C) Comparisons within the Near Eastern ancestry cluster

To determine the sex of the ancient individuals we computed the ratio of Y chromosome reads to the sum of chromosomes X and Y reads. As female individuals lack Y chromosome the ratio should be close to 0, whereas male individuals’ ratio should be significantly higher. We used a thresholding in this ratio of <0.03 for classifying a sample as female and >0.35 for male. Results of the determination can be found in **Additional Table 1.**

No apparent aneuploidies were identified in any of the samples neither in the autosomes nor on the sex chromosomes. Aneuploidies were studied by computing the mean coverage at 1240k SNPs of each chromosome divided by the mean coverage at 1240k SNPs of all autosomes, which should be around 1 for autosomes if no aneuploidies are present; 1 for females and 0.5 for males at the X chromosome if no aneuploidies are present, and 0.5 both at X and Y chromosomes for males if no aneuploidies are present. By comparing the coverage of sex chromosomes, we further added evidence for sex determination and classification of the newly reported samples (**Figure S6**).

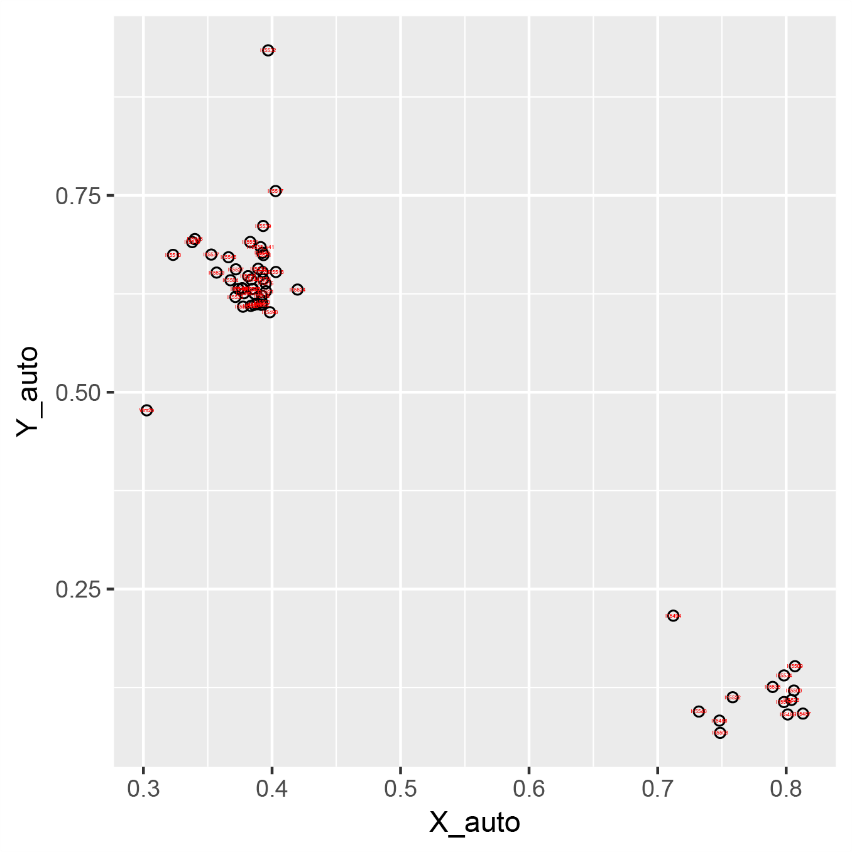

**Figure S6**. Scatterplot of Y chromosome and X chromosome mean coverage ratio for the newly reported ancient samples.

We assessed the possible presence of runs of homozygosity (ROHs) in the newly reported ancient samples by applying the method described in ^30^. No evidence of long ROHs was observed, indicating the absence of close kin unions in the ancestors of reported samples.

8. **Genome-wide analysis datasets**

To study the genetic ancestry of the newly sequenced individuals we built two datasets:

- The ‘HO’ dataset containing 6651 present-day individuals from worldwide populations ^1,2,31,32^, 37 newly reported present-day Serbs, and 44 newly reported present-day Greek individuals, all genotyped on the Human Origins Array, together with 3690 present-day individuals with whole-genome data from the SGDP, HGDP and 1000 genomes datasets ^2,31,33–38^, a set of 455 relevant ancient individuals with genome-wide data from previous publications ^11,13,31,39–56^ (Patterson et al. in submission), and our newly sequenced ancient individuals from Serbia. We kept 591,642 SNPs resulting from the intersection between the Human Origins array and the 1240k capture.

- The ‘1240k’ dataset contained the same individuals as the previously explained ‘HO’ dataset but excluded present-day individuals genotyped on the Human Origins array. We kept 1,054,671 autosomal SNPs, excluding SNPs of the 1240k array that are been specifically included based on their functional effects or located on the sex chromosomes.

In both datasets, for each individual we randomly sampled one allele at each SNP position to represent the data for that individual.

9. **Principal Component Analysis**

We performed Principal Component Analysis (PCA) on the HO dataset using the ‘smartpca’ program in EIGENSOFT (version 7.2.1). We projected ancient individuals onto the components computed on present-day individuals with “lsqproject:YES” and “shrinkmode:YES” ^57,58^.

We ran three different PCAs:

1. One for which PCs were computed using 1027 present-day West Eurasian individuals genotyped on the Human Origins array (**Figure 1**; **Figure S7**). Our newly reported individuals show a large spread on this West Eurasian PCA, with some individuals plotting very close to the Near Eastern cline, others very close to the European cline, and others in intermediate positions between these clines. Some ancient Serbian individuals plot on top of the present-day Balkan populations (including the newly reported present-day Serbian-speaking (n=37) individuals), who form a south to north cline in PC1, with southern populations such as Greeks plotting closer to present-day Near-Eastern populations, and northern populations plotting closer to other present-day Central/Eastern European populations.

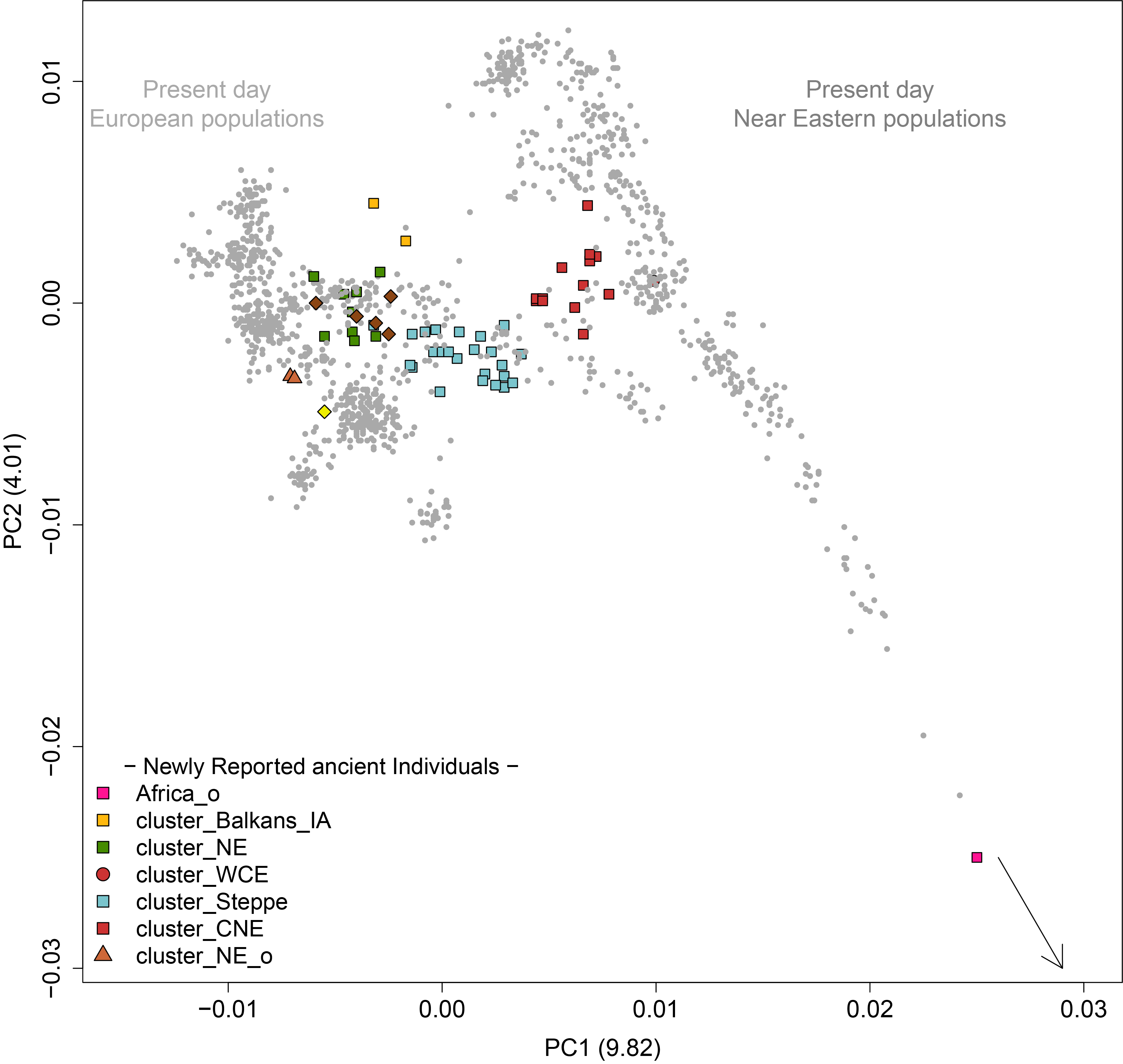

**Figure S7.** West Eurasian PCA with the newly reported ancient samples projected. This PCA is the zoom-out version of main text Figure 1 to fully visualize the West-Eurasian population structure.

2. One for which PCs were computed using 1226 present-day West Eurasian and African individuals genotyped on the Human Origins array (**Figure S8**). We use this PCA to investigate the genomic affinities of ancient individual I15499 from Viminacium, who plots outside West Eurasian genetic variation in the PCA.

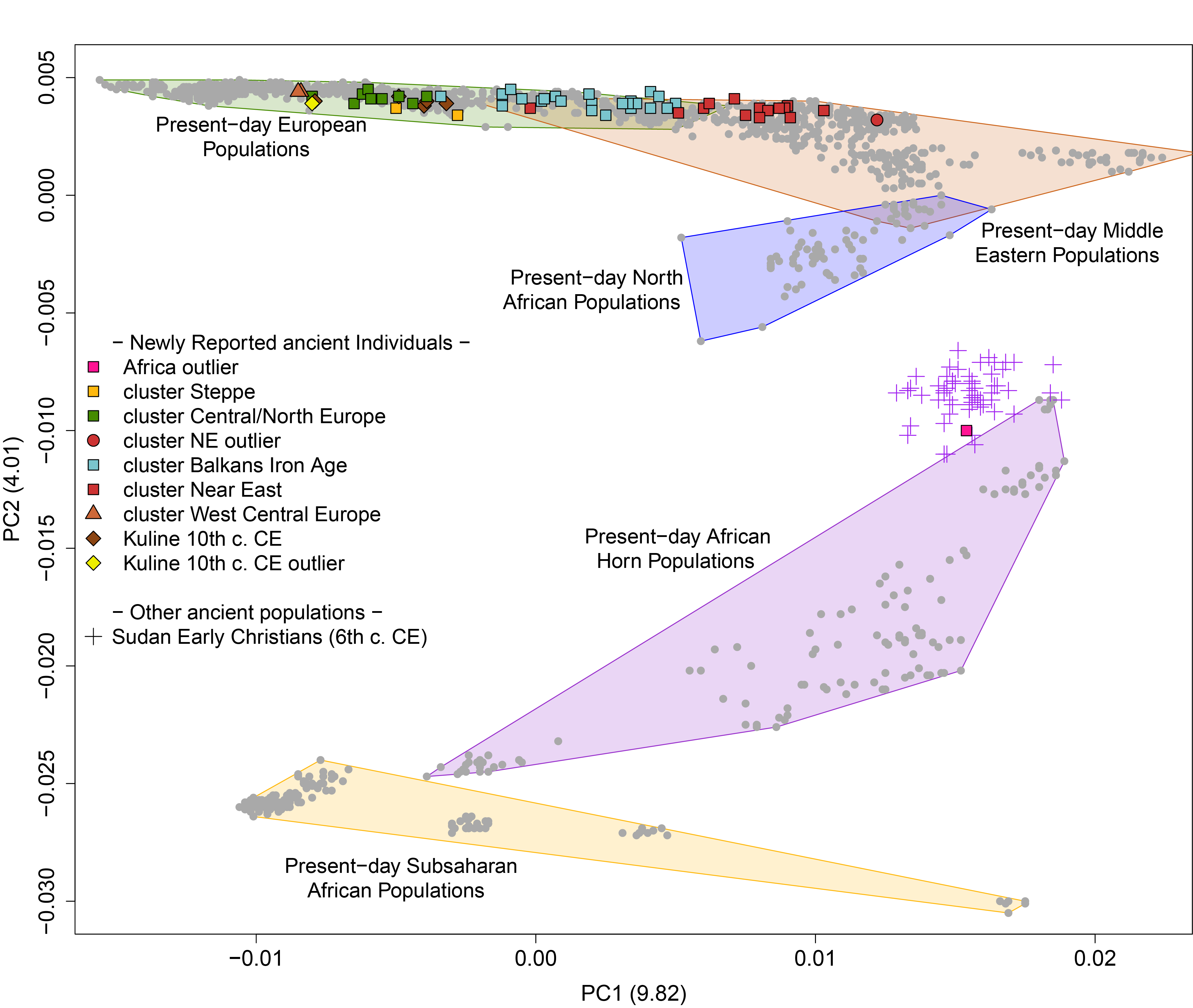

**Figure S8**. West-Eurasian and African PCA with the newly reported ancient individuals projected. The African outlier individual projected far outside of Western Eurasian genetic variation and close to present-day East African populations.

3. One for which PCs were computed using 407 present-day North-Europeans genotyped on the HO array (**Figure S9**). We designed this PCA to reveal more recent drift that could separate 3rd-6th centuries CE individuals from the 10th century CE individuals. These two groups of individuals yielded a similar position in the Western-Eurasian PCA (**Figure1; Figure S7**) but had significantly different ancestral origins when modelling using *qpWave/qpAdm* (Supplementary Section 11 & Supplementary Section 12). The design of this PCA was inspired by the Eurogenes blog:

(<https://eurogenes.blogspot.com/2017/10/tollense-valley-bronze-age-warriors.html>).

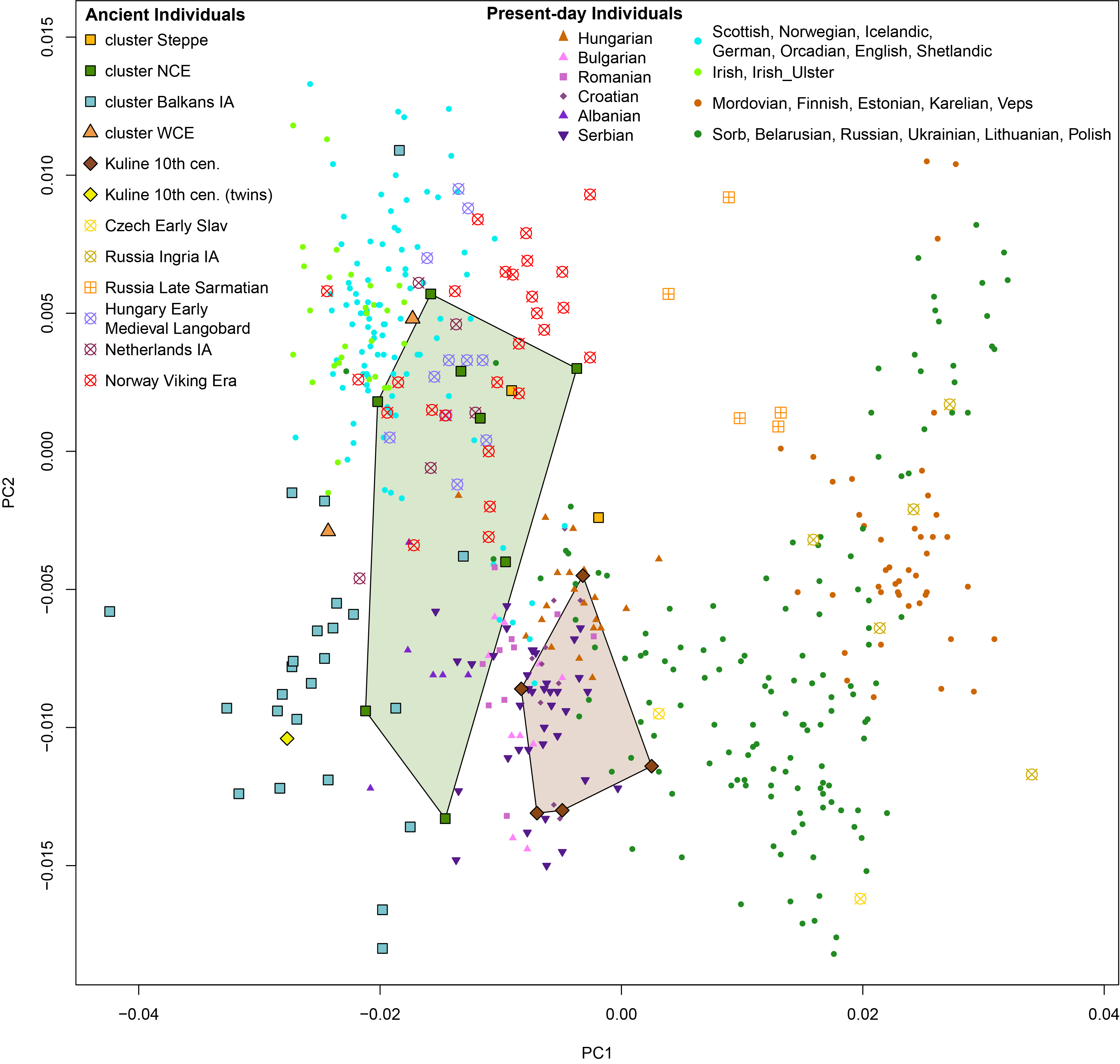

**Figure S9.** North European PCA with ancient samples projected.

10. **ADMIXTURE**

Model-based clustering analysis was performed using ADMIXTURE, version 1.3.0, ^59^ using the ‘HO’ reference dataset (Supplementary section 8), which included 2,562 modern individuals from worldwide populations, and the ancient individuals. SNPs in linkage disequilibrium were pruned from the dataset using PLINK, version 1.9 ^60^ with flags: –indep-pairwise 200 25 0.4, resulting in 308,524 SNPs. The ADMIXTURE program was run with cross-validation error flag (-cv) and *K* = 2 to *K* = 20, with 20 replicates for each value of K. Those replicates with the highest log-likelihood, and with an average pairwise similarity of over 0.98, were kept and plotted using *pong* version 1.4.1 ^61^. CV-error mean had a minimum at *K* = 20, although the lowest mean standard error was found in *K* = 18.

ADMIXTURE results (*K*=9) are consistent with the PCA in highlighting the diversity of the ancestral profiles of the Viminacum and Timacum Minus Slog individuals (**Figure S10**). *K*=9 is the lowest *K* for which components of ancestry related both to Iranian Neolithic farmers and European Mesolithic hunter-gatherers are maximized. The component maximized in Neolithic Iranians is the major component in the majority of the Roman Imperial period individuals, which could imply an increased influx of Near-Eastern or Eastern Mediterranean ancestry during the period of Roman control (pink in **Figure S10**). We also observe a general increased WHG related component in post-Roman samples from Timacum Minus Kuline Necropolis.

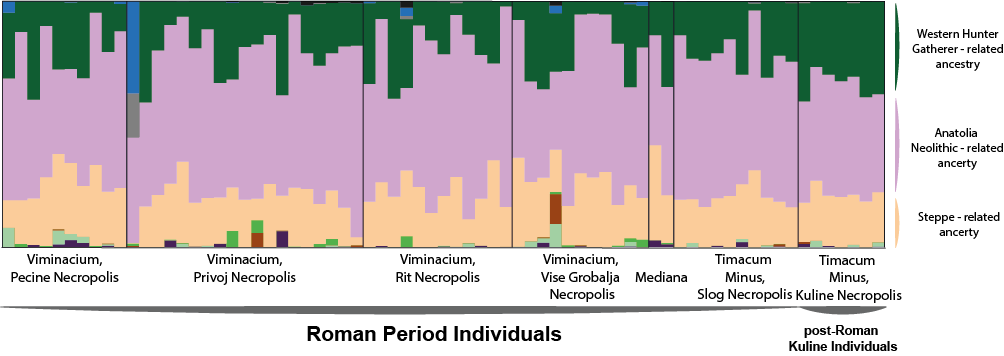

**Figure S10**. ADMIXTURE results (K=9) of the newly reported ancient samples. A) Roman period samples from Viminacum, Mediana and Timacum Minus. B) post- Roman samples from the 10^th^ Century Cal CE from Kuline necropolis in Timacum Minus. Maximized ancestral groups are shown in Green (Western Hunter Gatherer), Pink (Anatolia Neolithic), Orange (Steppe)

11. ***qpWave* tests for admixture**

The following sections explain the workflow for modelling the ancestry of the newly-reported ancient samples using the *f-*statistics framework on the ‘1240k dataset’ (Supplementary section 8).

Given the high ancestry heterogeneity observed in PCA and ADMIXTURE, we used the *qpWave* ^20^ from AdmixTools v.6. (https://github.com/DReichLab/AdmixTools) to group individuals with similar ancestry profiles into relatively homogenous clusters, whose ancestry we could then model using *qpAdm*. By increasing sample size for each modeled cluster, we increase the power to reject non-fitting models, as compared to an approach where samples are modelled individually.

Using the set of outgroups shown in **Table ST2**, we first ran in qpWave all possible pairwise comparisons of high-quality individuals (>300,000 SNPs) from Viminacium, Mediana and Timacum Minus (Slog Necropolis), and all possible pairwise comparisons between high-quality individuals (>300,000 SNPs) from Timacum Minus (Kuline necropolis). We chose to run a separate analysis of Kuline due to a chronological gap of 400-500 years with the other necropolises.

**Table ST2**. Set of outgroups used for qpWave and qpAdm. The IDs of the corresponding individuals of each population can be found in Additional File Table 3. (Abbreviations: EHG: Eastern Hunter Gatherer; HG: Hunter Gatherer; N: Neolithic; MLBA: Middle Late Bronze Age; IA: Iron Age; EIA: Early Iron Age)

| **Right Populations** | **References** |
| --- | --- |
| West_Africa_ancient | ^52^  ^55^ |
| Sudan_EarlyChristian | ^51^ |
| Tianyuan | ^56^ |
| EHG | ^48^ |
| Iron_Gates_HG | ^13^ |
| Levant_N | ^31^ |
| Anatolia_N | ^48^ |
| Iran_N | ^31^ |
| Yamnaya_Samara | ^48^ |
| Alalakh_MLBA | ^53^ |
| Iberia_IA | ^14^ |
| Mycenaean | ^46^ |
| Slovenia_EIA | _Patterson_ *_et al._*_, in submission_ |
| Netherlands_IA | _Patterson_ *_et al._*_, in submission_ |
| Russia_Ingria_IA | ^50^ |
| Steppe_IA | ^54^ |

The P-values of all pairwise *qpWave* tests are depicted is **Figure S11**. We observe 3 differentiated clusters that we named the Central/Northern European-related cluster (“CNE”), the Balkans Iron Age-related cluster (“Balkans IA”) and the Near Eastern-related cluster (“NE”) due to their position in PCA projections. Within each of these clusters, *qpWave* tests yielded high P-values indicating that individuals were symmetrically related to the populations in the outgroup set. Two individuals (I15527 and I15551) could not be assigned to any of the three large clusters, and were kept as single-individual clusters named Near Eastern outlier (NE outlier; I15551) and West/Central European-related clusters (NWE cluster; I15527).

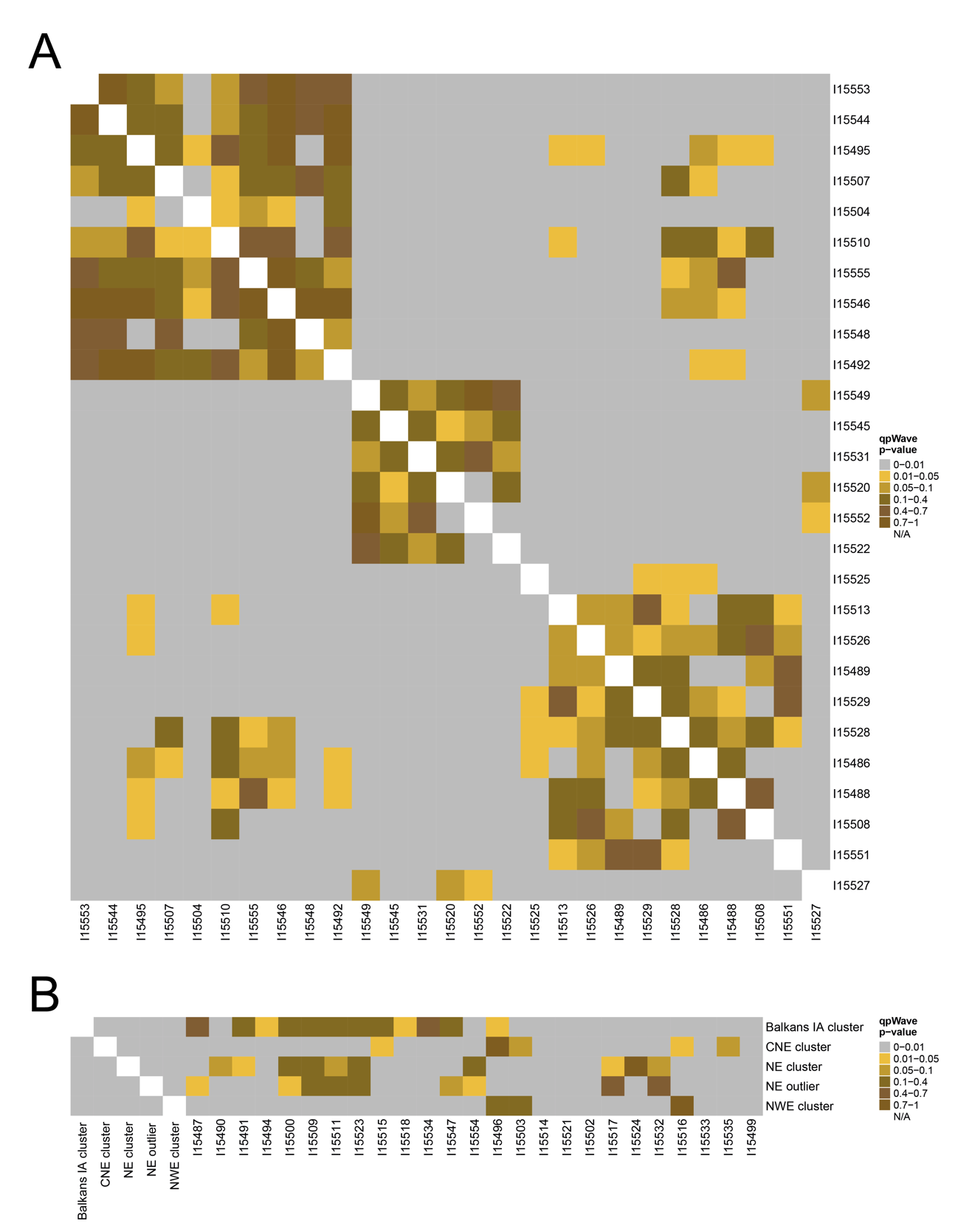

**Figure S11.** qpWave clustering of newly reported ancient samples. A) Comparisons among high coverage individuals (> 300,000 SNPs). B) Comparisons of lower coverage individuals (15,000 to 300,000 SNPs) with clusters.

For Kuline, all individuals were consistent with forming a clade, with the exception of the pair of twins.

Next, we tested in *qpWave* the lower coverage individuals with <300,000 SNPs (excluding those with fewer than 15,000 SNPs) against the five clusters identified in the first step (**Figure S11.B**) and that were either assigned to one of these five clusters or to new clusters (Africa outlier and Steppe-related cluster) if they were not consistent with forming a clade with any of them. For borderline cases, we also considered PCA positions for aiding cluster assignment. Cluster assignment for each individual is shown in **Table ST3**. Finally, clusters defined in the two previous steps were tested against each other to check that they were differentially relative to the outgroup. None of the cluster significantly shared ancestry against the used set of outgroups.

**Table ST3.** Final assignment of ancient samples to shared ancestry clusters based on qpWave tests.

| **ID** | **Cluster** |
| --- | --- |
| I15499 | Africa outlier |
| I15533 | Steppe cluster |
| I15535 | Steppe cluster |
| I15553 | Balkans IA cluster |
| I15554 | Balkans IA cluster |
| I15544 | Balkans IA cluster |
| I15490 | Balkans IA cluster |
| I15495 | Balkans IA cluster |
| I15507 | Balkans IA cluster |
| I15504 | Balkans IA cluster |
| I15518 | Balkans IA cluster |
| I15510 | Balkans IA cluster |
| I15555 | Balkans IA cluster |
| I15547 | Balkans IA cluster |
| I15546 | Balkans IA cluster |
| I15548 | Balkans IA cluster |
| I15492 | Balkans IA cluster |
| I15515 | Balkans IA cluster |
| I15534 | Balkans IA cluster |
| I15494 | Balkans IA cluster |
| I15491 | Balkans IA cluster |
| I15511 | Balkans IA cluster |
| I15487 | Balkans IA cluster |
| I15500 | Balkans IA cluster |
| I15523 | Balkans IA cluster |
| I15509 | Balkans IA cluster |
| I15521 | Balkans IA cluster |
| I15549 | CNE cluster |
| I15545 | CNE cluster |
| I15531 | CNE cluster |
| I15520 | CNE cluster |
| I15552 | CNE cluster |
| I15514 | CNE cluster |
| I15496 | CNE cluster |
| I15503 | CNE cluster |
| I15522 | CNE cluster |
| I15525 | NE cluster |
| I15513 | NE cluster |
| I15502 | NE cluster |
| I15526 | NE cluster |
| I15489 | NE cluster |
| I15524 | NE cluster |
| I15529 | NE cluster |
| I15517 | NE cluster |
| I15532 | NE cluster |
| I15550 | NE cluster |
| I15528 | NE cluster |
| I15486 | NE cluster |
| I15488 | NE cluster |
| I15508 | NE cluster |
| I15551 | NE outlier |
| I15516 | NWE cluster |
| I15527 | NWE cluster |

12. ***qpAdm* admixture modelling**

In *qpWave* (Supplementary Section 11) we grouped individuals from Viminacium, Timacum Minus and Mediana into clusters. In the following section, we attempt to model the ancestry of each of these relatively homogenous clusters.

We used the same set of outgroups as for the *qpWave* tests (**Table ST2**): *West_Africa_ancient, Sudan_EarlyChristian, Tianyuan, EHG, Iron_Gates_HG, Levant_N, Anatolia_N, Iran_N, Yamnaya_Samara, Alalakh_MLBA, Iberia_IA, Mycenaean, Slovenia_EIA, Netherlands_IA, Russia_Ingria_IA, Steppe_IA,* unless otherwise justified for specific tests. This set of populations includes groups distantly related (both geographically and chronologically) to our Serbian ancient individuals, and more proximate groups that could form a clade with the true sources of ancestry in them.

We tested all possible 1-way, 2-way and 3-way combinations of populations in our outgroup population set, using them as sources and leaving the remaining populations as outgroups in the model. We then checked whether a 1-way model was sufficient to explain the ancestry in the *test* population. If no 1-way model showed a good fit (p-value > 0.05), we looked for plausible 2-way models, and, if the 2-way model was still not fitting, we looked for 3-way models. We computed standard errors for *qpAdm* tests using a weighted block jackknife over 5-Mb blocks.

12.1 *Balkans Iron Age cluster*

No one-way model fits the ancestry of this group. However, three 2-way models work:

$$0.916 \left( \pm0.01 \right)Mycenaean+ 0.084 (\pm0.01) Iron\_Gates\_HG$$

[P-value = 0.031];

$$0.798 (\pm0.018) Mycenaean + 0.202 (\pm0.018) Netherlands\_IA$$

[P-value = 0.056];

$$0.665 (\pm0.027) Mycenaean + 0.335 (\pm0.027) Slovenia\_EIA$$

[P-value = 0.095]

All three models feature a southern Balkans Bronze Age population, the Mycenaean group, and a group falling on or beyond the European cline in PCA. The first model fails when Villabruna is included in the outgroup set, indicating that the ancestry not explained by the Mycenaean group cannot be derived solely from a pre-Neolithic European group (such a model is also implausible chronologically). The model with highest P-value, a mixture between Mycenaean-related ancestry and Slovenia Early Iron Age-related ancestry, is also the most plausible chronologically and geographically, and is concordant with the evidence from the PCA, as this cluster of individuals falls within the cline delimited by the Bronze Age (e.g. Mycenaeans) and Iron Age-Roman Period (e.g. Aegean-like individuals from the Greek colony of Empúries) Aegeans on the right and Croatia Middle Bronze Age-Iron Age and Slovenia Early Iron Age on the left. Thus, we interpret this group as the descendants of Balkans Iron Age populations, representing the indigenous component present at Viminacium and Timacum during Roman Imperial times. Two radiocarbon dates were obtained from this cluster yielding 129-247 calCE (1835±20 BP, PSUAMS-8560) and 261-418 calCE (1685±20 BP, PSUAMS-8725), indicating that individuals who were direct descendants of Balkans IA populations, although eventually disappeared through admixture with later incoming groups (they are not found in present-day Balkans), did survive until at least the fourth century CE [**Table ST1**].

12.2 *Near Eastern cluster*

There are no 1-way models nor 2-way models fitting the ancestry of this group. The only model that worked (P-value = 0.096) using this set of outgroups is:

$$0.90 \left( \pm0.022 \right)\mathrm{Mycenenan} + 0.082 (\pm0.015) Iran\_N + 0.019 (\pm0.017) Netherlands\_IA$$

This model suggests that the ancestry in this group deviates from Mycenaean-related ancestry in the direction of Iran Neolithic-related ancestry, but it does not provide good information about the proximal ancestry of this cluster. Furthermore, the ancestry proportion assigned to Netherlands_IA is very low (barely different from 0), and the 2-way model featuring Iran_N and Mycenaean with Netherlands_IA in the outgroups fails with P-value=0.007, likely indicating that to model this cluster of individuals one needs more central or northern European-related ancestry than that included in the model Iran_N+Mycenaean, but not as much as that present in Netherlands_IA. A plausible reason no other models work is that, even though this group clearly has a shift towards Anatolian Late Chalcolithic-Early Bronze Age groups in PCA, only one post-Neolithic Near Eastern group is included in the initial outgroup set, the Alalakh Middle-Late Bronze age group from Northern Levant, which is not a good proxy fom the ancestry in this cluster. Therefore, we tested 1-way models featuring an Anatolian Chalcolithic (LC) or Early Bronze Age (EBA) population as the only source (from West to East: *Anatolia_Barcin_LC*, *Anatolia_Gondurle_EBA*, *Anatolia_CamlibelTarlasi_LC*, *Anatolia_Ikiztepe_LC*, *Anatolia_Arslantepe_LC*, *Anatolia_Arslantepe_EBA*). Using the full set of outgroups in **Table ST2**, we obtained the results depicted in **Table ST4**.

**Table ST4.** Results of one-way modeling of the NE cluster with ancient Anatolian populations as sources (Abbreviations: LC;Late Chalcolithic; EBA:Early Bronze Age)

| **Source** | **P-value** |
| --- | --- |
| Anatolia_Barcin_LC | 0.047 |
| Anatolia_Gondurle_EBA | 2.02E-17 |
| Anatolia_CamlibelTarlasi_LC | 7.62E-44 |
| Anatolia_Ikiztepe_LC | 1.18E-18 |
| Anatolia_Arslantepe_LC | 2.36E-55 |
| Anatolia_Arslantepe_EBA | 5.64E-34 |

Only the Late Chalcolithic individual from Barcın (near the Sea of Marmara) was a plausible proxy for the ancestry in this Near Eastern-related cluster of Roman-period Serbian individuals, although chronologically it is thousands of years earlier. To improve the fit of these models and based on the findings that Chalcolithic and Bronze Age populations from Anatolia had different levels of Iran Neolithic-related ancestry ^53^, we ran the same models again but added Iran Neolithic as a source together with the Anatolian groups, to account for possible differences in Iran Neolithic-related ancestry between the available Anatolian populations and the true source of ancestry in our test population. We obtained:

$$0.934 \left( \pm0.029 \right) Anatolia\_Barcin\_LC + 0.066 (\pm0.029) Iran\_N$$

[P-value = 0.109];

$$0.883 \left( \pm0.020 \right) Anatolia\_Gondurle\_EBA + 0.117 (\pm0.020) Iran\_N$$

[P-value = 1.41E-11];

$$0.926 \left( \pm0.020 \right) Anatolia\_CamlibelTarlasi\_LC + 0.074 (\pm0.020) Iran\_N$$

[P-value = 5.45E-40];

$$0.947 \left( \pm0.020 \right) Anatolia\_Ikiztepe\_LC + 0.053 (\pm0.020) Iran\_N$$

[P-value = 5.22E-15];

$$0.993 \left( \pm0.020 \right) Anatolia\_Arslantepe\_LC + 0.007 (\pm0.020) Iran\_N$$

[P-value = 8.57E-52];

$$1.018 \left( \pm0.023 \right) Anatolia\_Arslantepe\_EBA + -0.018 (\pm0.023) Iran\_N$$

[P-value = 9.41E-31];

We confirm that adding a small proportion of Iran Neolithic-related ancestry provides a good fit with *Anatolia_Barcin_LC* as the main ancestry source, while the other Anatolian sources continue to provide poor fits. Given that *Anatolia_Barcin_LC* is represented by a single individual with ~576K SNPs, we checked whether the good fit for this group could be due to a reduced power to reject poorly-fitting models, as compared to the other Anatolian groups that include several individuals ^62^. We thus tested the same model of Anatolian group + Iran_N but using single individuals with similar numbers of available SNPs as *Anatolia_Barcin_LC* [**Table ST5**]:

**Table ST5**. Two-Way NE cluster modeling with an ancient Anatolian and Iran Neolithic as sources. (Abbreviations: LC;Late Chalcolithic; EBA:Early Bronze Age)

| **Source 1** | **SNPs** | **Proportion**  **Source 1** | **Proportion**  **Iran_N** | **SE** | **P-value** |
| --- | --- | --- | --- | --- | --- |
| Anatolia_Arslantepe_LC_  ART038 | 319571 | 0.980 | 0.020 | 0.029 | 3.27E-08 |
| Anatolia_Arslantepe_LC_  ART023 | 353091 | 1.013 | -0.013 | 0.029 | 2.03E-11 |
| Anatolia_Gondurle_EBA_  I2499 | 382206 | 0.938 | 0.062 | 0.029 | 6.72E-10 |
| Anatolia_Arslantepe_LC_  ART019 | 452291 | 1.085 | -0.085 | 0.035 | 2.48E-15 |
| Anatolia_Arslantepe_LC_  ART005 | 471618 | 0.939 | 0.061 | 0.029 | 1.46E-11 |
| Anatolia_CamlibelTarlasi_LC_  CBT005 | 509734 | 0.945 | 0.055 | 0.028 | 1.65E-12 |
| Anatolia_CamlibelTarlasi_LC_  CBT010 | 529018 | 0.942 | 0.058 | 0.029 | 3.51E-09 |
| Anatolia_CamlibelTarlasi_LC_  CBT001 | 609436 | 0.917 | 0.083 | 0.030 | 1.26E-12 |
| Anatolia_CamlibelTarlasi_LC_  CBT004 | 615196 | 0.979 | 0.021 | 0.032 | 3.63E-07 |
| Anatolia_Arslantepe_EBA_  ART011 | 640615 | 1.054 | -0.054 | 0.036 | 1.30E-09 |

In all cases, even when using individuals with substantially fewer SNPs than the *Anatolia_Barcin_LC* individual, models are strongly rejected. This confirms that the well-fitting model for *Anatolia_Barcin_LC* is not a consequence of limited power to reject poorly-fitting models due to this group including only one individual.

To understand why *Anatolia_Barcin_LC* + *Iran_N* provides a good fit but models featuring other Anatolian groups + *Iran_N* are strongly rejected, we added as a third source the Balkans Iron Age-related cluster of Serbian Roman-period individuals modelled in the previous section (Supplementary section 12.1), to account for European-related ancestry that could be missing in the Anatolian groups providing poor fits, but present in *Anatolia_Barcin_LC* and our test population (the Near Eastern-related cluster of Roman-period Serbian individuals) (**Table ST6**).

**Table ST6**. Three-Way NE cluster modeling with an ancient Anatolian, Iran Neolithic, and Balkans Iron Age-related Roman-Serbian cluster as sources. (Abbreviations: LC;Late Chalcolithic; EBA:Early Bronze Age)

| **Source 1** | **Proportion**  **Source 1** | **SE** | **Proportion Iran_N** | **SE** | **Proportion Balkans_IA cluster** | **SE** | **P-value** |
| --- | --- | --- | --- | --- | --- | --- | --- |
| Anatolia_  Gondurle_EBA | 0.480 | 0.040 | 0.114 | 0.014 | 0.406 | 0.038 | 0.203 |
| Anatolia_  Camlibel  Tarlasi_LC | 0.420 | 0.032 | 0.097 | 0.013 | 0.483 | 0.030 | 0.002 |
| Anatolia_  Ikiztepe_LC | 0.527 | 0.042 | 0.074 | 0.014 | 0.399 | 0.038 | 0.020 |
| Anatolia_  Arslantepe_LC | 0.449 | 0.031 | 0.057 | 0.013 | 0.493 | 0.026 | 0.197 |
| Anatolia_  Arslantepe_EBA | 0.449 | 0.034 | 0.056 | 0.015 | 0.495 | 0.028 | 0.391 |

The fit greatly improved by including the Balkans IA cluster as a source, suggesting that these Anatolian populations are missing a European/Balkan ancestry component that is present in *Anatolia_Barcin_LC*. and in the Near Eastern-related cluster of Serbian individuals which is also chronologically more plausible.

To sum up, the individuals with Near Eastern affinities in Viminacium (the Near Eastern cluster) can be modelled as deriving the vast majority of their ancestry ultimately from groups related to Late Chalcolithic western Anatolians, although with slightly more Iran-Neolithic-related ancestry than the samples with currently available data. Even when including the modeling Indigenous Balkan populations, they must still derive roughly half their ancestry from groups related to Anatolians related to those from the Chalcolithic and Bronze Age. In the future, genomic data from Hellenistic and Roman Anatolia will provide more proximate sources for the ancestry of this cluster.

12.3 *Near Eastern outlier (I15551)*

This individual dated to 242-375 calCE (1750±20 BP, PSUAMS-8561) is the only one from Timacum Minus with Near Eastern genomic affinity, falling right onto the Near East cline in PCA. For the same set of outgroups, a one-way model works having as a source *Alalakh_MLBA* (P-value = 0.3588). When we tested one-way models using the same Anatolian groups as for the main Near Eastern cluster (Supplementary section 12.2), we found good fits for *Anatolia_Arslantepe_LC* and *Anatolia_Arslantepe_EBA* [**Table ST7**]. We interpret these results as evidence for this individual having Eastern Anatolian/Northern Levant-related ancestry.

**Table ST7**. Results of one-way modeling of the NE outlier I15551 with ancient Anatolian populations as sources. (Abbreviations: LC;Late Chalcolithic; EBA:Early Bronze Age)

| **Source** | **P-value** |
| --- | --- |
| Anatolia_Barcin_LC | 0.00014 |
| Anatolia_Gondurle_EBA | 2.53E-05 |
| Anatolia_CamlibelTarlasi_LC | 8.03E-07 |
| Anatolia_Ikiztepe_LC | 0.048 |
| Anatolia_Arslantepe_LC | 0.273 |
| Anatolia_Arslantepe_EBA | 0.338 |

12.4 *African outlier (I15499)*

Only one 1-way model worked for this individual (P-value = 0.134), having ancestry related to 8th-9th century individuals from Northern Sudan (*Sudan_EarlyChristian)* (Sirak et al., in review). This model points to a clear Northeastern African origin of this individual, consistent with the historically and archaeologically well-documented interactions between the Early Roman Empire (**Table ST1**) with its African territories and foreign kingdoms such as Nubia or Meroe ^63^

12.5 *North-Western European cluster*

Only a single 1-way model featuring *Netherlands_IA* (P-value = 0.119) fits the ancestry of this group composed of two Viminacium individuals. Based on their PCA position, we tested whether Iron Age populations from Northeastern France (*France_GrandEst_IA2*) (Patterson et al., in review) could be a good surrogate of their ancestry. Indeed, we obtained a well-fitting model (P-value=0.864). These results point to a North-Western European origin for these two individuals, which is supported by the information provided by uniparental markers as one of the individuals harbored the R1b-U106 Y-chromosome haplogroup, found at high frequencies in Germanic-speaking countries today and at very low frequencies in the Balkans ^64^.

12.6 *Central/Northern European cluster*

No 1-way or 2-way models fitted the ancestry of these individuals. All the well-fitting 3-way models have *Netherlands_IA* as one of the sources, suggesting the presence of northern European-related ancestry. Given that these individuals date to ~250-550 cal CE (**Table ST1**), it is plausible that they also harbour ancestry related to the local cluster of ancient Serbian individuals modelled in supplementary section 12.1 (*Balkans Iron Age cluster*), some of whom date slightly earlier (129-247 calCE (1835±20 BP, PSUAMS-8560)) while others are contemporaneous (261-418 calCE (1685±20 BP, PSUAMS-8725)) to this *Central/Northern European cluster*. Therefore, we tested a 2-way model with *Balkans Iron Age cluster* + *Netherlands_IA*, keeping all the outgroups except *Netherlands_IA*:

$$0.349\left( \pm0.032 \right) Balkans\_Iron\_Age\_cluster+ 0.651 \left( \pm0.032 \right)Netherlands\_IA$$

[P-value = 0.00014];

The model does not provide a good fit, so we looked for groups that could represent a good proxy for the unmodelled ancestry in this group. Given that two contemporaneous individuals from Viminacium (modelled in section 12.7) are shifted in PCA towards Steppe populations from the first half of the first millennium CE, we hypothesized that the individuals from the *Central/Northern European cluster* also harbored some ancestry related to these steppe groups. We thus add *Russia_Late_Sarmatian* ^65^ from the Eastern Pontic-Caspian steppe to the previous model:

$$0.383 \left( \pm0.028 \right) Balkans Iron Age cluster$$

$$+ 0.481 \left( \pm0.045 \right)Netherlands\_IA$$

$$+ 0.136 \left( \pm0.030 \right)Russia\_Late\_Sarmatian$$

[P-value = 0.039];

The model significantly improves with 13% ancestry related to *Russia_Late_Sarmatians*. This indicates that this cluster of Serbian individuals have ancestry related to the local Balkans substratum, ancestry related to Central/Northern European populations, and ancestry related to contemporaneous nomadic steppe groups. As the current proxy for Central/Northern European-related ancestry in the model is *Netherlands_IA*, separated by ~300-900 years and ~1600 kilometers from our test individuals, we wanted to test whether other groups with similar ancestry but more proximate both in time and space could also be a good proxy for this type of ancestry. We therefore repeated the previous model but substituted *Netherlands_IA* with a group of individuals from a Langobard-associated cemetery in Hungary displaying Central/Northern European ancestry ^40^; IDs: SZ12, SZ16, SZ24, SZ30, SZ41, SZ7, SZ9), who are roughly contemporaneous to our individuals of interest and separated by ~350 kilometers:

$$0.356 \left( \pm0.027 \right) Balkans Iron Age cluster$$

$$+ 0.502 \left( \pm0.043 \right) Hungary\_Langobards$$

$$+ 0.142 \left( \pm0.029 \right) Russia\_Late\_Sarmatian$$

[P-value = 0.198]

This model again provides a good fit, with almost identical ancestry proportions as compared to the previous model including *Netherlands_IA*. Thus, these results point to gene flow from Central/Northern Europe into the Roman Empire Balkans territories starting at leaset in the late third century CE, consistent with the known arrival of Germanic groups beginning during the Crisis of the Third Century ^66^.

In order to effectively model this cluster of individuals, a steppe population carrying Asian-related ancestry was required, plausibly meaning that Central/Northern Europe groups interacted with steppe populations in Eastern Europe before reaching the Balkans and admixing with the local Balkans substratum.

12.7 *Steppe cluster*

No 1-way model fit the ancestry of this pair of individuals from Viminacium, one of them directly dated to 246-365 calCE (1745±15 BP, PSUAMS-8591). Two 2-way models worked using the set of outgroups:

$$0.390 (\pm0.025) Anatolian\_N + 0.610 (\pm0.025) Yamnaya\_Samara$$

[P-value = 0.054];

$$0.637 (\pm0.027) Mycenean + 0.363 (\pm0.027) Yamnaya\_Samara$$

[P-value = 0.582]

Both these models feature a group high in Anatolia Neolithic-related ancestry (either *Anatolia_N* itself or the *Mycenaean* group), and *Yamnaya_Samara*, suggesting that this cluster carries high levels of both ancestry derived from European Neolithic populations (abundant in the Balkans) and steppe-related ancestry. However, these models are too distal for our Roman-period groups and do not provide precise information about their proximal ancestral origins, likely because we lack more proximate sources of ancestry in the outgroups list.

Based on their position in PCA, and following a similar approach as for the contemporaneous *Central/Northern European cluster* in the previous section (supplementary section 12.6), we tested a model including the local Balkans substratum represented by the *Balkans Iron Age cluster* (supplementary section 12.1), and *Russia_Late_Sarmatian* as a proxy for groups living in the Pontic-Caspian steppe during the first half of the first-millennium CE. We kept the full the set of outgroups, and, unlike the previous section did not include a Central/Northern European source to check whether this cluster could model without this type of ancestry:

$$0.434 \left( \pm0.036 \right)Balkans\_Iron\_Age\_Cluster$$

$$+ 0.566 \left( \pm0.036 \right)Russia\_Late\_Sarmatian$$

[P-value = 0.516]

We confirm that this model fits well the ancestry of this pair individuals, who do not need Central/Northern European-related ancestry that is required by the cluster in the previous section. These results again demonstrate gene-flow into the Balkans from the Steppe during the third-fourth centuries CE, but with a different mix of ancestry sources. An extensive sampling of both the Balkans and the Steppe during this time period will further clarify the precise sources and the demographic impact of this event.

12.8 10th-century individuals from Kuline necropolis (Timacum Minus)

In this section, we attempt to model the ancestry of the main cluster of individuals from Kuline necropolis, dated to the 10th century CE (897-1021 calCE (1075±15 BP, PSUAMS-8555)), while the twins from Kuline are modelled in the next section (supplementary section 12.9). In PCA, this cluster of 10th century individuals plot on top of the *Central/Northern European cluster* modelled in supplementary section 12.6, who date to more than 400 years earlier. Based on this evidence, we tried the same model of *Balkans Iron Age cluster* + *Hungary_Langobards* + *Russia_Late_Sarmatian*, that provided a good fit for the *Central/Northern European cluster*:

$$0.436 \left( \pm0.036 \right) Balkans\_Iron\_Age\_cluster$$

$+0.510 \left( \pm0.055 \right)Hungary$_Langobards

$$+0.054 (\pm0.032) Russia\_Late\_Sarmatian$$

[P-value = 1.18E-05];

The model does not provide a good fit for the ancestry in this group, implying that the similar position in the West Eurasian PCA is a projection artifact masking a subtly different ancestry makeup.

The North European PCA (**Figure S5**) is driven by more recent drift and shows a clear separation between both groups, with individuals in the *Central/Northern European cluster* closer to present-day Germanic-speaking groups, and the 10th century individuals from Kuline closer to present-day Slavic-speaking groups from Eastern Europe, and on top of present-day Serbs.

We thus tested a 2-way model having the local Balkans ancestry substratum (*Balkans Iron Age cluster*) as one source and a proxy for Northeastern European ancestry as a second source:

$$0.562 \left( \pm0.023 \right)Balkans Iron Age cluster$$

$$+ 0.438 \left( \pm0.023 \right)Russia\_Igria\_IA$$

[P-value = 0.0302]

When we use Iron Age individuals from Ingria ^50^ as the second source, the method provides a reasonable fit, with P-value = 0.03 and roughly equal proportions of *Balkans Iron Age cluster*-related ancestry and *Russia_Ingria_IA*-related ancestry. Substituting *Russia_Ingria_IA* by two individuals from Brandysek (Czech Republic) dated to 660-774 calCE sequenced in 1240k ^39,48^ did not provide a good fit (P-value 0.01). The lack of available genomic data from populations living in Eastern Europe during the first millennium CE prevented us from using more proximate sources, both temporally and geographically, for the Eastern European-related ancestry in this group.

Our interpretation of these results is that these 10th century CE individuals from Kuline necropolis (Timacum Minus) already carried Northeastern European ancestry related to Slavic-speaking groups known to have settled in the region in the 7^th^ century ^66^ as well as local Balkans-related ancestry indicating that these incoming groups extensively mixed with the people they encountered. The demographic impact of this event in the Balkans is further explored in Supplementary section 13 using present-day Balkan populations.

We used qpAdm to attempt to detect sex bias in this group of individuals by modeling the X chromosome using the same Slavic-proxy and local cluster are sources ^67^. This model also fitted the X chromosome ancestry, although with a higher standard error:

$$0.235 \left( \pm0.1 \right)Balkans Iron Age cluster$$

$$+ 0.765 \left( \pm0.1 \right)Russia\_Igria\_IA$$

[P-value = 0.113]

The X-chromosome model significantly deviates from the autosome model (Z-score = 3.187) showing a higher proportion the *Russia_Ingria_IA*-related ancestry in this chromosome. This result implies that immigrant females had significantly a greater impact on the resulting mixed populations of the Balkans, suggesting a bigger female to male ratio migrated during the Slavic people movement and settlement in the Balkans. However, we have not detected a significant sex bias in present-day Balkan populations present in the Simons Genome Diversity Project ^35^. The extent of this sex bias is yet to be further explored in other post-Slavic migration and present-day Balkan populations.

12.9 Twins from Kuline, Timacum Minus (I15538 and I15539)

In this section we model the ancestry of two 10th century CE individuals (892-989 calCE (1115±15 BP, PSUAMS-8592)) from Kuline identified as identical twins (Supplementary Section 7) and who displayed a different ancestry makeup compared to the other individuals from the same necropolis. Data from both individuals were merged to achieve higher power in *qpAdm* analysis. We found that only a single 1-way model with *Slovenia_EIA* provided a good fit (P-value = 0.067) for these individuals. Trying other 1-way models featuring groups not present in the outgroup set, and keeping the whole set as outgroups, we found that *Croatia_MBA_EIA* ^13^ (P-value = 0.355), *France_SouthEast_IA2* (Patterson *et al.*, in review) (P-value = 0.12928549) and *France_Occitanie_IA2* ^42^(p-value = 0.069) also yielded a good fit for their ancestry. This means that different groups from Southern Europe can correctly model the ancestry of these twins, all geographically located at either East or West of the Alps. We argue that an origin West of the Alps is more likely because a) Populations with ancestry similar to *Slovenia_EIA or Croatia_MBA_EIA* likely did not exist anymore in the Balkans during the 10th century CE, and b) their mtDNA lineage, H1e1a6, was initially discovered in Iberian present-day populations ^68^ and has only been found in five other ancient individuals so far: I19987 and I19989 from Iron Age Iberia (Patterson *et al.*, in review), I3579 from Early Medieval Iberia ^14^, an early 20^th^ century CE individual from Iberia ^69^, and MS10585 from 5^th^ century BCE Sardinia ^70^. This suggests an Iberian or more generally a southwestern European origin for this lineage, and a more plausible ancestral origin in southwestern European for these identical twins.

13 ***qpAdm* admixture modeling of present-day Balkan populations**

Based on the finding of Northeastern-related ancestry in the 10th century individuals from Kuline, likely associated to the Slavic migration during the Early Medieval period, we sought to study whether this signal persisted in present-day Balkan and Aegean populations, which would imply a long-term demographic impact of this event in the region. The following steps and tests were carried out using the ´HO´ dataset, which includes 591,642 SNPs (Supplementary section 8). We used the same set of outgroups as explained in Supplementary section 11 (**Table ST2**): *West_Africa_ancient, Sudan_EarlyChristian, Tianyuan, EHG, Iron_Gates_HG, Levant_N, Anatolia_N, Iran_N, Yamnaya_Samara, Alalakh_MLBA, Iberia_IA, Mycenaean, Slovenia_EIA, Netherlands_IA, Russia_Ingria_IA, Steppe_IA,* unless otherwise stated.

First, we attempted to model the ancestry of present-day Balkan populations as one-way models with different groups whose ancestry derived entirely from pre-Roman Balkans populations: *Croatia_MBA_EIA* (Mathieson *et al.,* 2018), the cluster of Roman-period individuals from Timacum Minus and Viminacium modelled in supplementary section 12.1 (*Balkans Iron Age cluster*)*,* and 400 BCE - 200 CE individuals from the Greek colony of Empúries (Spain) with fully Aegean-related ancestry (*Greek_Empuries* ^14^) (**Table ST8**). These three groups acted as representatives of northern, central and southern Balkans-related ancestry, respectively. If the 1-way models provided a good fit to the data, this would indicate genetic continuity in the Balkans since prior to the Roman period and no significant long-term demographic impact of the Slavic migration or other population movements in the region over the past ~2,000 years. However, all the models failed with extremely low P-values, strongly rejecting population continuity in the Balkans since pre-Roman times, and documenting a history of mixture.

**Table ST8**. Present-day Balkan (and surrounding) populations one-way modeling

|  | **p-value** | | |
| --- | --- | --- | --- |
| **Balkan (and surrounding)**  **Present-day populations** | **Balkans Iron Age**  **cluster** | **Croatia_**  **MBA_EIA** | **Greek_**  **Empuries** |
| Albanian | 2.378E-32 | 2.19E-12 | 1.68E-31 |
| Bulgarian | 1.292E-76 | 4.13E-17 | 2.96E-59 |
| Cretan | 3.913E-44 | 3.09E-44 | 3.57E-15 |
| Croatian | 1.694E-170 | 2.28E-31 | 3.62E-100 |
| Greek_Cyclades | 9.427E-25 | 2.08E-31 | 7.10E-13 |
| Greek_Dodecanese | 2.615E-61 | 5.37E-64 | 2.53E-13 |
| Greek_Macedonia | 3.456E-50 | 5.89E-18 | 6.14E-35 |
| Greek_Peloponnese | 3.816E-24 | 8.13E-20 | 1.93E-21 |
| Hungarian | 2.285E-275 | 1.42E-48 | 1.32E-132 |
| Romanian | 6.512E-98 | 2.16E-17 | 6.34E-69 |
| Kuline 10th c. CE | 9.162E-47 | 1.19E-11 | 1.30E-52 |
| Serbian | 5.190E-183 | 1.20E-24 | 3.97E-95 |

We then tried to model the present-day groups as a two-way model. Similar to the modelling of 10^th^ century individuals from Kuline (supplementary section 12.1), we try models with one local Balkans source (either *Balkans Iron Age cluster*, *Croatia_MBA_EIA* or *Greek_Empuries*), and a proxy for Northeastern European-related ancestry (either *Russia_Ingria_IA* or present-day populations from Eastern Europe).

**Table ST9**. Present-day Balkan populations two-way modeling with Balkans Iron Age cluster and Russia_Ingria_IA as sources.

| **Balkan**  **Present-day**  **populations** | ***Balkans Iron Age***  ***cluster*** | **SD** | ***Russia_Ingria_IA*** | **SD** | **p-value** |
| --- | --- | --- | --- | --- | --- |
| Albanian | 0.753 | 0.022 | 0.247 | 0.022 | 2.98E-03 |
| Bulgarian | 0.622 | 0.021 | 0.378 | 0.021 | 7.10E-02 |
| Cretan | 1.004 | 0.034 | -0.004 | 0.034 | 0 |
| Croatian | 0.458 | 0.019 | 0.542 | 0.019 | 0.153449 |
| Greek_Cyclades | 0.989 | 0.027 | 0.011 | 0.027 | 1.67E-14 |
| Greek_Dodecanese | 1.211 | 0.039 | -0.211 | 0.039 | 0 |
| Greek_Macedonia | 0.763 | 0.019 | 0.237 | 0.019 | 2.28E-10 |
| Greek_Peloponnese | 0.866 | 0.024 | 0.134 | 0.024 | 1.16E-11 |
| Hungarian | 0.372 | 0.018 | 0.628 | 0.018 | 0.07882 |
| Romanian | 0.567 | 0.020 | 0.433 | 0.020 | 2.925E-4 |
| Kuline 10th c. CE | 0.569 | 0.026 | 0.431 | 0.026 | 0.174897 |
| Serbian | 0.514 | 0.017 | 0.486 | 0.017 | 0.014691 |

A model having *Balkans Iron Age cluster* (as the local source) and *Russian_Ingria_IA* (as the Northeastern European-related source) fitted for three present-day Balkan populations, Hungary, Croatian and Serbian with P-value>0.01 (**Table ST9**), as well as for the Kuline 10th c. CE with almost identical mixture proportions as in the 1240k dataset (supplementary section 12.1).

However, this model did not fit the ancestry of the remaining more southern (except Romanian) populations, who instead required a more local source represented by *Greek_Empuries*, and present-day Mordovian or Russian as proxy for Northeastern European-related ancestry. These models fit the ancestry for the remaining Balkans populations (**Table ST10**; **Table ST11**), with ~30-55% Northeastern European-related ancestry.

**Table ST10**. Present-day Balkan two-way modeling with *Greek_Empuries* and present-day Mordovians as sources.

| **Balkan**  **Present-day**  **populations** | ***Greek_Empuries*** | **SD** | **Mordovian** | **SD** | **p- value** |
| --- | --- | --- | --- | --- | --- |
| Albanian | 0.624 | 0.021 | 0.376 | 0.021 | 0.388 |
| Romanian | 0.471 | 0.017 | 0.529 | 0.017 | 0.175 |
| Bulgarian | 0.523 | 0.017 | 0.477 | 0.017 | 0.309 |
| Greek_Cyclades | 0.804 | 0.024 | 0.196 | 0.024 | 0.00025 |
| Greek_Dodecanese | 0.926 | 0.027 | 0.074 | 0.027 | 1.97E-12 |
| Greek_Macedonia | 0.641 | 0.018 | 0.359 | 0.018 | 0.203 |
| Greek_Peloponnese | 0.713 | 0.021 | 0.287 | 0.021 | 0.048 |

**Table ST11**. Present-day Balkan two-way modeling with Greek_Empuries and present-day Russians as sources.

| **Balkan**  **Present-day**  **populations** | ***Greek_Empuries*** | **SD** | ***Russian*** | **SD** | **p- value** |
| --- | --- | --- | --- | --- | --- |
| Albanian | 0.624 | 0.021 | 0.376 | 0.021 | 0.203 |
| Romanian | 0.451 | 0.017 | 0.549 | 0.017 | 0.685 |
| Bulgarian | 0.507 | 0.017 | 0.493 | 0.017 | 0.497 |
| Greek_Cyclades | 0.811 | 0.024 | 0.189 | 0.024 | 2.49E-05 |
| Greek_Dodecanese | 0.943 | 0.028 | 0.057 | 0.028 | 4.87E-13 |
| Greek_Macedonia | 0.633 | 0.018 | 0.367 | 0.018 | 0.098 |
| Greek_Peloponnese | 0.710 | 0.021 | 0.290 | 0.021 | 0.015 |

These two-way models failed to model the ancestry in Aegean islander populations (*Greek_Cyclades*, *Greek_Dodecanese* and *Cretan*), which displayed a very low P-value (**Table ST10**; **Table ST11**). Based on the shift of this cline towards the Levant, observed by their higher PC1 values in the West-Eurasian PCA (**Figure 1**, **Figure S7**), we tested if changing the local source population to one that yield Near-Eastern Related ancestry would achieve a fitting model. We used I7833 individual as source who had a significant Near-Eastern signal and high coverage (Lazaridis et al., in submission), from Roman Greece previous to the Slavic migrations (252-412 cal CE), who we refer as *Roman_Greek*. Indeed, replacing *Greek_Empuries* by this individual resulted in a good-fitting model for the Aegean islander population. We also tried to model the Cypriots, who occupy the most “southern” extreme of this Greek cline.

Based on the clear shift towards present-day Cypriot that these groups presented in PCA, we test whether adding present-day Cypriot as a proxy for Near Eastern related ancestry could improve the fit. (**Table ST12**; **Table ST13**) Indeed, adding this third source resulted in a good-fitting model for the three groups, who required a substantial amount of

**Table ST12.** Present-day Greek Islander two-way model with individual I7833 as local source and Mordovians as Slavic proxy.

| **Greek Islanders** | ***Roman_Greek*** | | **Mordovian** | **SD** | **pvalue** |
| --- | --- | --- | --- | --- | --- |
| Greek_Cyclades | | 0.846 | 0.154 | 0.04 | 0.686963 |
| Greek_Dodecanese | | 0.985 | 0.015 | 0.047 | 0.677534 |
| Cretan | | 0.864 | 0.136 | 0.041 | 0.683756 |
| Cypriot | | 1.073 | -0.073 | 0.049 | 0.923474 |

**Table ST13.** Present-day Greek Islander two-way model with individual I7833 as local source *and* Russians as Slavic proxy.

| **Greek Islanders** | ***Roman_Greek*** | **Russian** | **SD** | **pvalue** |
| --- | --- | --- | --- | --- |
| Greek_Cyclades | 0.837 | 0.163 | 0.04 | 0.779971 |
| Greek_Dodecanese | 0.989 | 0.011 | 0.047 | 0.668616 |
| Cretan | 0.862 | 0.138 | 0.041 | 0.675598 |
| Cypriot | 1.077 | -0.077 | 0.049 | 0.933824 |

We observe Northeastern Europe-related ancestry in the Cyclades and Crete which are more closely located to the Greek mainland. This ancestry signal (absent in Iron Age and Roman Balkan populations) decreases from North to South in the Balkans, but it is still substantial in populations from these Aegean islands. However, this North-Eastern signal is not significant in the farther islands: the Dodecanese and Cyprus, who even rejects the model by having negative values in the former.
